# A survey of the mentor-mentee relationship in early career research (ECR): Implications for publishing and career advancement in the STEMM disciplines

**DOI:** 10.1101/2024.10.29.620999

**Authors:** Ronan Lordan, Michael Wride, Íde O’Sullivan

## Abstract

Early career researchers (ECRs) are the most abundant workforce in the fields of science, technology, engineering, mathematics, and medicine (STEMM). ECRs are generally mentored by experienced principal investigators (PIs) who direct the research objectives. The ECR mentee-mentor partnership can be mutually beneficial, but it is a critical relationship for ECRs with implications for publishing and career development. In this study, a mixed methods approach involving a survey, X polls (formally Twitter), and semi-structured interviews were used to determine how the ECR mentor-mentee relationship affects ECRs and their perceptions of career development in STEMM. To address this aim, two main research questions were developed: (1) Who do the STEMM ECR mentees and mentors currently perceive to be ECRs? and (2) What characteristics of the ECR mentee-mentor relationship are perceived to be important for ECR publishing practices and career progression? The main survey findings indicate that respondents largely from the life sciences and health field and they consider Ph.D. researchers and postdoctoral researchers as ECRs, but mentors also to a greater extent than mentees perceive new PIs (< 2 years experiences) as ECRs. Survey respondents also mostly agree on publishing practices regarding journal selection criteria such as journal scope and impact factor, although mentees appear to favor journal prestige and open access publishing more than mentors. This is likely due to the importance of visibility early in one’s career. Mentees take the lead in preparing manuscripts for submission, although a minority have described issues regarding authorship disputes. Finally, setting clear expectations, being collegial, mutually respectful, and having regular communication was identified by survey respondents and interviewees as integral features of a healthy ECR mentee-mentor relationship. In conclusion, the mentee-mentor relationship is critically important for ECR career development, and the findings of this study have wider implications for the development of effective ECR mentee and mentor training programs across the STEMM disciplines.

## 1.0. Introduction

The fields of science, technology, engineering, mathematics, and medicine (STEMM) are critically important to the advancement of society, economic growth, and educating future generations to deal with impending global challenges [1]. Early career researchers (ECRs) are the most abundant and most diverse cohort of scientists in STEMM [2] who make up the bulk of the scientific workforce [3]. ECRs are critical drivers of innovation within STEMM and important contributors to the improvement of science [4, 5].

Despite the multiple definitions that exist, there is no universally accepted definition of what is an ECR, sometimes also referred to as an early-stage researcher (ESR) [6]. This is because definitions may differ depending on the views of different countries, academic cultures, or practices, or indeed stipulations of hiring committees, grant, or funding agencies [3, 6, 7]. ECRs are commonly described by universities, government bodies, and funders as non-tenured researchers, who either have received their doctorate and are currently in a research position or have been in research positions but are currently doing a doctorate. This may include graduate and medical students, postdoctoral researchers, young clinical investigators, and some definitions even include newly appointed independent investigators beginning their career as principal investigators (PIs) [3, 6, 7]. However, these definitions are not unilaterally accepted and can vary from country to country, or depending on the academic department or funding agencies [6].

Discrepancies in defining ECRs has wider implications for ECR identity, intellectual belonging, and career outcomes for ECRs in the highly competitive arena of STEMM [8, 9], including exclusion from funding opportunities such as the NIH 4 year maximum time post Ph.D. to be considered eligible for K99/R00 pathway to independence awards [10]. This type of exclusionary practice based on years post Ph.D. fails to take into account the complexity or time it takes to complete a postdoc in a field studying aging or longitundinal research, disadvantaging ECRs in some branches of science.

Researchers in STEMM assess successes by the so-called traditional metrics that include the number of papers published, journal impact factors, total citations, H-index, and the amount of funding obtained. These metrics are used for the purpose of hiring, promotion, tenure, and to provide future resources and rewards by academic departments and/or funding agencies [6, 11–13]. However, they can also be assessed for the purpose of hiring scientists outside of academia in industry or governmental roles [14]. While it is well-known that these metrics do not adequately reflect a researcher’s ‘success’ [13, 15–18], ECRs who are interested in pursuing a career in academia are dependent on a favorable publication record [9, 19–21]. This has led to a ‘publish or perish’ academic culture in a stagnating hiring environment where there are now 6.3 Ph.D. graduates for every biomedical tenure track position in the United States [22]. Indeed, metrics show that of the 56,000 postdocs in the United States, only 15–20% of them will become tenure-track faculty [23, 24], despite 80% of postdoctoral researchers indicating their preference for a tenured career in academia [25]. Of those lucky enough to gain professorships, the majority tend to be male [26]. These competitive conditions have led to an exodus of postdoctoral researchers out of academic career paths in STEMM [22, 27], due to economic challenges, poor compensation, gender bias, and in general the “publish or perish” culture of academia making postdoctoral researchers feeling undervalued [28–30]. Consequently, postdoctoral researchers are recognized as a highly vulnerable group [31].

ECRs are largely reliant on the mentorship, supervision, and training by more experienced principal investigators who direct the objectives of a research group, and who therefore influence the research outputs and publications of the ECR, affeting their career trajectory [32]. The degree to which ECRs contribute to decisions surrounding publication strategies, which directly impacts their career trajectory [6, 33], is not clear and likely depends on the academic culture within a field. Good mentorship is critically important to ECRs as it is a time of personal and professional growth [9] and mentee-mentor relations substantially impact the experience and productivity of ECRs [34].

There is a dearth of literature relating to the ECR mentee-mentor relationship and how it may affect publishing practices among ECR mentees, mentors, and academics in general. Qureshi, Gröschner [35] developed a mentoring program and focused on patterns of communication between mentees and mentors and its impact on career development using semi-structured interviews. They reported positive improvement in communication, but longitudinal studies are required to determine effects on career outcomes. Nicholas, Rodríguez-Bravo [6] conducted an interview-based study of 116 ECRs regarding publishing and authorship practices to determine whether ECRs were more likely to adopt modern approaches to publishing such as open access publishing and social media versus the more traditional approaches of their mentors in aiming for high impact established publishers. They found that ECRs were more likely to adhere to convention due to the precarious employment environment that relies on the traditional approaches to publishing to evaluate candidate. Other investigators have focused on ECR wellbeing. A well conducted survey of over 650 Australian ECRs reported that a number of factors including job insecurity, workplace culture, questionable research practices, and mentorship were impacting job satisfaction despite a clear love of science among the responders [36]. The aim of our study was to specifically target the role of the ECR mentee-mentor relationship and what constraints exist interpersonally or institutionally that affect the publishing practices and career progression of ECRs.

In this study, social media was leveraged to recruit ECR mentees and ECR mentors to take part in Twitter polls (now known as X), surveys, and semi-structured interviews to determine how the ECR mentee-mentor relationship affects ECRs and their perceptions of publishing practices and career development in the STEMM disciplines. To address the overarching aim, two main research questions were developed: (1) Who do the STEMM disciplines (ECR mentees and mentors) currently perceive to be ECRs; (2) What characteristics of the ECR mentor and ECR mentee relationship are perceived to be important for ECR publishing practices and career development.

## 2.0 Methods

In this mixed-methods research project three approaches were utilized. Quantitative data was obtained using a survey and Twitter polls. Qualitative data was obtained using open-ended questions in the survey and a semi-structured interview with ECRs and ECR mentors. Ethical review and approval were granted by the Faculty of Arts, Humanities, and Social Sciences, at the University of Limerick, Ireland (Approval Number: 2022-02-10-AHSS). The Twitter polls were conducted from March 8^th^ to March 15^th^, 2023. Surveys and interviews were conducted from March 20^th^ to April 4^th^, 2023. Participants of the survey and interview provided informed consent to take part in this study. Due to a lack of a clear definition for an ECR, the invitation to participate in the study was open to anyone who identified as either an ECR mentee or mentor.

## 2.1. Survey

An invitation to take part in the survey was distributed on social media platforms including Discord, Facebook, LinkedIn, Slack, and Twitter in March 2023. These social media platforms were chosen due to the known presence of academic communities. For example, on Slack the “Future PI Slack” channel has greater than 5,000 users. Indeed, Twitter has a vast academic community that has a track record of being used for research [37, 38] and the distribution of surveys [39, 40]. Hashtags such as #AcademicChatter and #EarlyCareerResearcherChat were used to distribute the invitations on the social media platforms. The survey was split into three sections. Section 1 was a common section that was designed to gather demographic information and to ask common questions to both ECR mentors and mentees. These questions sought information regarding age, research discipline, institution type, and country origin. The survey then branched into either an ECR mentee section (Section 2) or an ECR mentor section (Section 3) depending on whether the respondent identified as an ECR mentee or mentor. The questions in these sections of the survey addressed similar topics regarding the relationship between ECR mentees and mentors, their writing and publication strategies, and the career development of ECRs. The survey was developed using questions adapted from published research on similar topics [6, 36] and additional questions were created to address our research needs. The survey questions were piloted by ECR mentors (n=2) and ECR mentees (n=2) prior to distribution. Quantitative data was gathered from Likert scale type questions. Qualitative data was gathered using open-ended questions, from which responses were analyzed using word clouds and word association analyses (Nicholson et al. 2022). Respondents were free to skip questions they were uncomfortable answering.

### 2.2. X poll analyses

X, formerly known as Twitter, is a microblogging platform on social media with approximately 350 million active users. X was a popular place for academics where collaborations and research can occur between individuals from all stages of their careers, nationally and internationally. This has been exemplified in studies that have used X for research initiatives regarding the pandemic [41], hashtag analyses of various campaigns in the biomedical field [38], research of the natural products [37], and various other projects in health research [42]. Prior to the transition from Twitter to X, reports published by Nature estimated that 13% of researchers are regular users of the platform [43]. There has since been a reduction of users due to the change in leadership [44, 45]. However, X continues to be a platform that scientist communicate on and use for research in various fields [46, 47]. Therefore, X users were asked to interact with four polls regarding ECRs and publishing relating to topics asked in the surveys using the main author’s X account. The polls were simultaneously placed on Twitter March 8^th^, 2023, and were circulated for 7 days on Twitter. Links guiding participants to the Twitter polls were also circulated on other social media platforms including Discord, Slack, and Facebook. Twitter polls are limited in their content to 280 characters and only 4 responses (25 characters for each option). The four polls were designed under the constraints of the Twitter character and response limits to emulate similar questions that were asked in the survey to get a broader academic view on these important topics (Table 1). The first poll question (Poll 1) was asked to gain an insight regarding what the wider academic Twitter community consider to be an ECR. Poll 2 was designed to address the perception of which ECR outputs (e.g., number of publications, citation counts, high impact factor papers, or varied skill set) are considered more important by the academic Twitter community for career development and hiring purposes in academia. Poll 3 was asked to determine what factors affect journal selection for publishing manuscripts among the Twitter academic community. Finally, poll 4 asks if the academic Twitter community have ever experienced author disputes. All Twitter poll responses were anonymous.

**Table 1:**
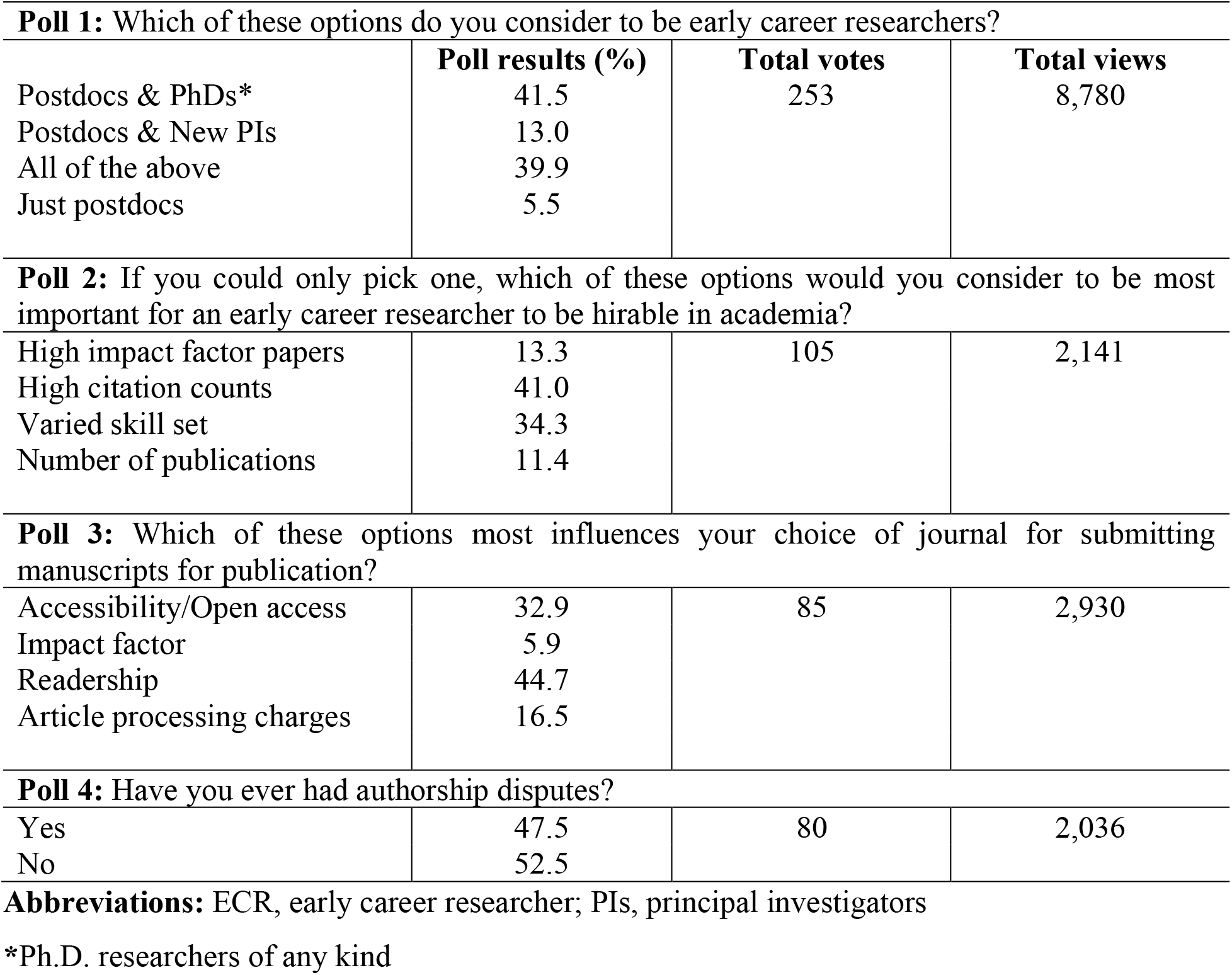
Four polls were deployed on Twitter to obtain additional data regarding the definition of an ECR from a broad academic perspective. Twitter users were asked about what factors academics perceive to be important for job prospects in academia, and publishing practices such as journal selection and authorship disputes.

### 2.3. Semi-structured interviews

Those who completed the survey were presented with an option to provide their informed consent and contact details to take part in a semi-structured interview for approximately 30 minutes. Within 7 days of responding to the survey, participants were invited to take part in the interview via email. In total, 5 ECR mentees and ECR mentors 3 were recruited. The interviews took place online using Microsoft Teams. A set of similar questions were posed to both the ECR mentor and mentees that were designed to allow the respondents to provide their opinions on topics relating to ECR mentorship, career progression, and publication practices. The interviews were recorded and transcribed using Microsoft Teams. The quality of transcription was checked against the recordings by the author. Once the raw transcription data was anonymized the recordings were deleted to preserve anonymity. For this study, quotes that exemplify the findings of the interviews have been rereferred to in this manuscript (Supplementary Tables S9 and S10). However, a full thematic analysis of these interview transcripts is the subject of another ongoing study intended for publication. The questions asked to both mentors and mentees are listed in Table S11.

### 2.4. Data presentation and analyses

Quantitative data was presented as bar charts, pie charts, and dot plots using GraphPad Prism 9 (GraphPad Software, Boston, MA, USA). Percentages were calculated per number of respondents to each individual question. Word clouds, also known as tag clouds or term clouds, were generated using a free word cloud generator (https://www.freewordcloudgenerator.com/generatewordcloud, Salt Lake City, UT, USA) to present an overview of common words used among written responses. Word clouds provide a graphical representation of text data through differences in text size and color that provide the viewer with a rapid and intuitive sense of the text [48, 49]. In this study, the larger and greener the text, the more frequent the word was used by respondents. Interview transcripts were produced using the Microsoft Teams transcription feature, which was then manually assessed for accuracy. All transcripts were deidentified and anonymized. Excerpts from the transcripts of both mentors (n =3) and mentees (n = 5) were quoted and presented in text and placed in table format throughout the manuscript.

## 3.0. Results and Discussion

### 3.1. Early career researcher mentor and mentee survey respondent demographics

A total of 138 survey responses were recorded of which 106 were ECR mentees and 28 were mentors. The majority of the participants of the survey were female (56%; Figure 1A), in their early thirties (Figure 1B), white race (Figure 1D), and ECR mentees (Figure 1C). The survey captured respondents from over 30 countries, the majority of which currently worked in either Australia, Ireland, the United Kingdom, or the United States (Figure 1E). Most participants indicated that they were either working in research at a university (36.6%), teaching and researching at a university (31.3%) or a student (19.4%; Figure 1G). The majority of respondents indicated that they were either a postdoctoral researcher (36.6%) or a postgraduate student (28.4%), with a minority of participants identifying as an assistant professor/professor (17.2%; Figure 1G). Indeed, most respondents indicated that they worked in the fields of biological sciences (45.5%) or medical and health sciences (18.7%; Figure 1H). The vast majority of respondents (92%) indicated that they had received a doctorate degree (Ph.D.), with many also indicating that they had achieved a master’s degree (56%).

**Figure 1:**
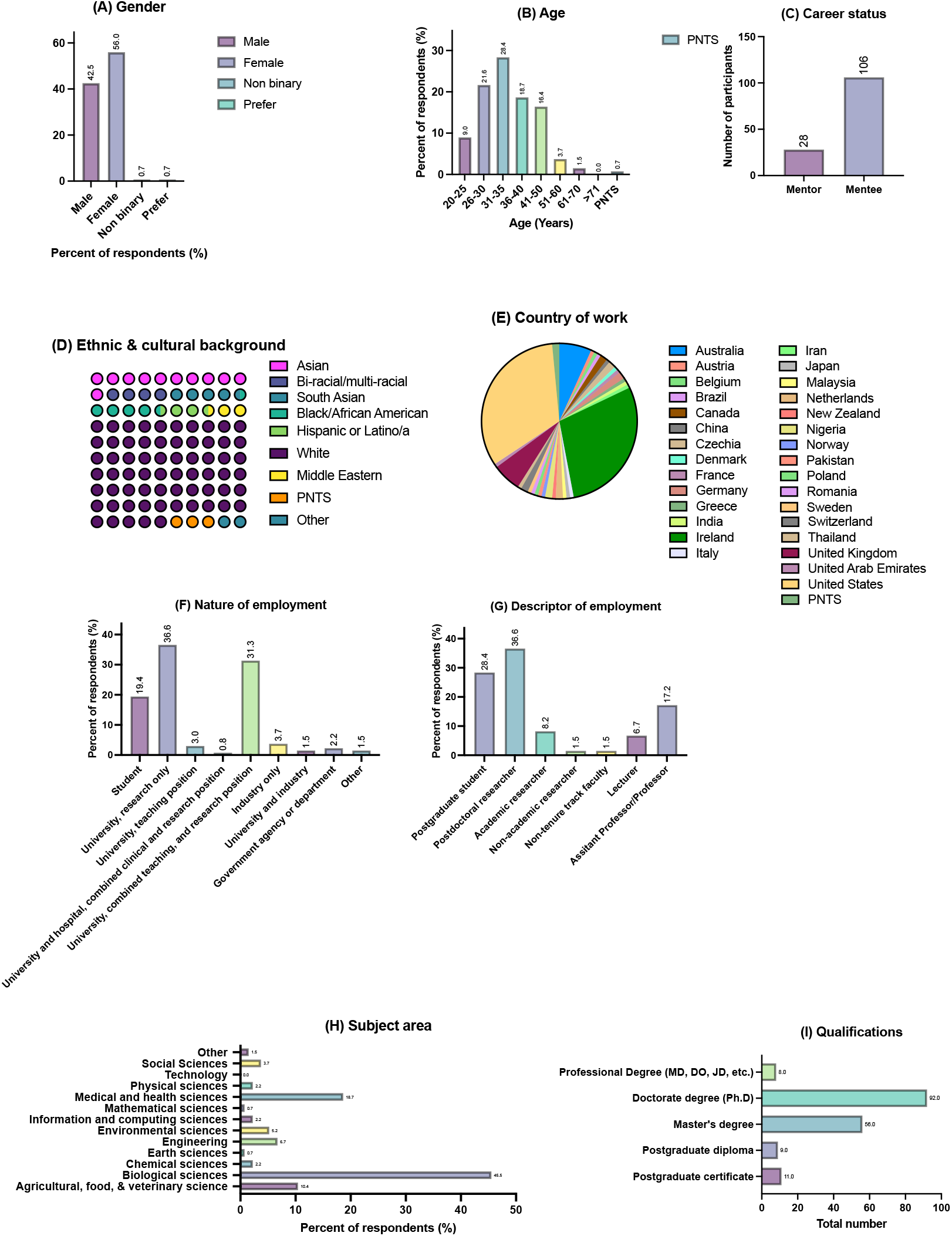
The demographic data of the survey participants are presented in panels A-I. (A) The gender of the respondents (n = 134). (B) The age of the respondents (n = 134). (C) The number of mentors (n = 28) and mentees (n = 106) who took part in the study. (D) The percent distribution of different ethnicities and cultural backgrounds of the participants (n =131, n = 3 PNTS). (E) The percent distribution of the current country of work of each respondent (n = 132, n = 2 PNTS). (F) The nature of employment of each participant expressed as a percent of the total respondents (n = 112). (G) The descriptor of employment of each participant expressed as a percent of the total respondents (n = 134). (H) The general subject area of the respondents expressed as a percent of the total respondents (n = 134). (I) The number and type of degrees held by respondents (excluding a bachelor’s degree) (n = 134). Abbreviation: PNTS, prefer not to say.

### 3.2. Defining early career researchers (ECRs)

To uncover current perceptions regarding defining ECRs in the STEMM fields, both STEMM ECR mentees and mentors were surveyed, and X polls were conducted among the online academic community. Our findings indicate that there is a broad range of perceptions regarding the definition of an ECR among the STEMM community (Figure 2 and Table 1). The majority of ECR mentee respondents selected that Ph.D. researchers (72.6%) and postdoctoral researchers either 0-2 years active (64.2%), 0-5 years active (61.3%) or any postdoctoral researchers (51.9%) were considered ECRs. The majority of ECR mentors considered Ph.D. researchers (85.7%) and postdoctoral researchers 0-2 years active (85.7%) as ECRs. Notably mentors indicated that principal investigators with less than 2 years’ experience (67.9%) should be considered ECRs, whereas a lesser number of mentees agreed (48.1%).

**Figure 2:**
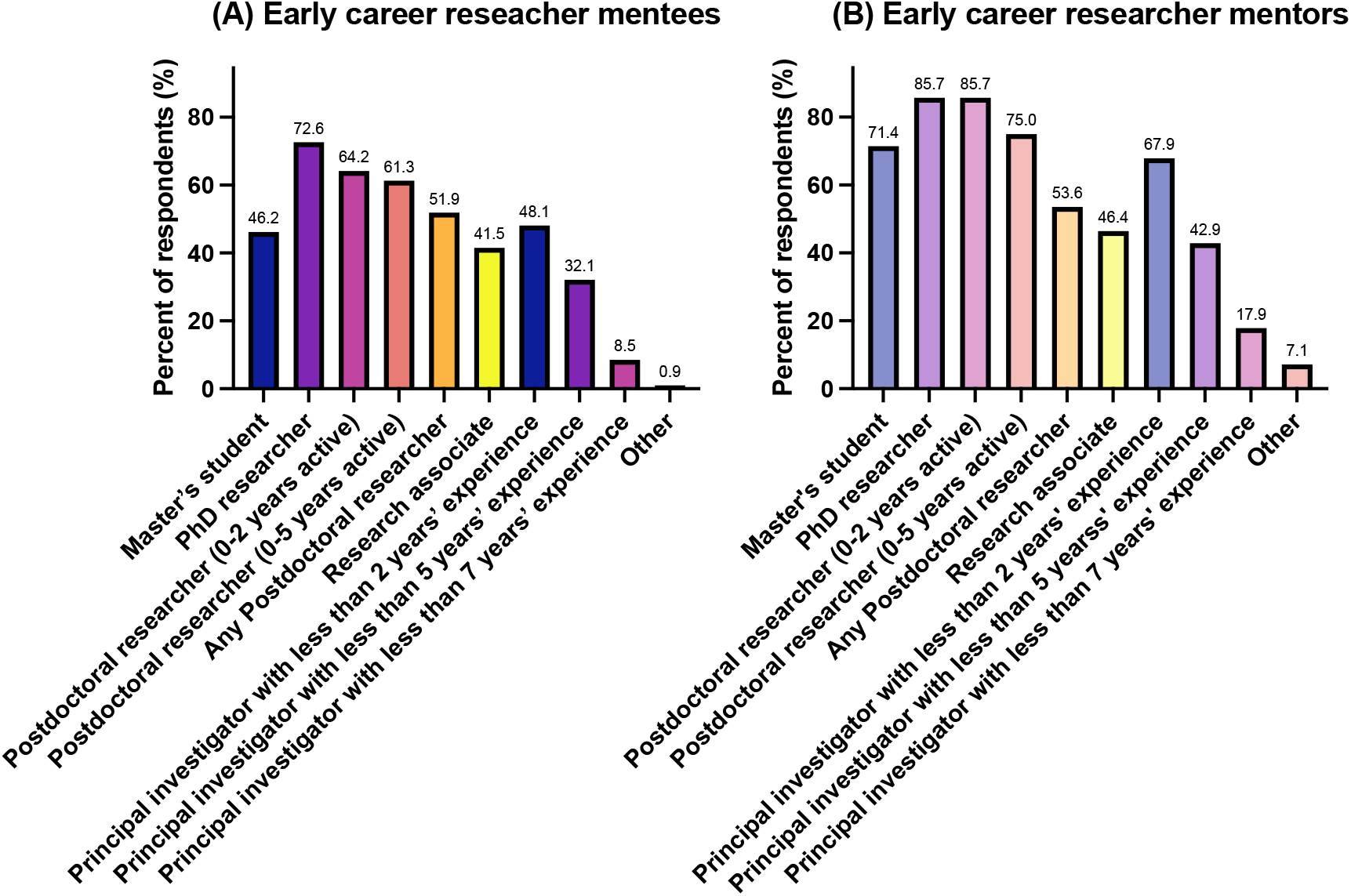
ECR mentees and mentors were asked to select which options they considered to be ECRs. A total of 106 mentees and 28 mentors responded. A percentage was calculated from the total number of mentees or mentors who selected each option.

A poll was also devised to determine who academics on Twitter considered to be ECRs (Table 1). Respondents could only choose one of these four options. Out of a total of 253 votes obtained from 8,780 views of the poll (2.9% response rate), the majority considered the collective option of “postdocs and PhDs” as ECRs (41.5%), whereas a subset of respondents did not consider Ph.D. researchers as ECRs as they selected “Just postdocs” (5.5.%) or “Postdocs & PIs” (13%) as ECRs. The remaining respondents (39.9%) viewed Ph.D. researchers, postdocs, and new PIs as ECRs as they selected “All of the above”. These findings generally corroborate the survey data presented in Figure 2.

Collectively, the findings of the survey and Twitter polls confirm that Ph.D. researchers and postdoctoral researchers are the majority fraction of the academic work force who are considered ECRs according to the STEMM ECR mentee and mentor respondents in this study. A considerable number of ECR mentees and mentors also recognized master’s candidates, research associates, and PIs with 2-5 years’ experience as ECRs. These findings further emphasize the varied perceptions of ECR identity that exist, likely driven by the multitude of definitions that exist in the literature and adopted by funding agencies. Therefore, it is important that academic departments and funding agencies who stratify candidates based on one’s academic identity consider inclusive criteria for hiring, promotion, tenure, resource allocation, and/or rewards if ECRs are to be considered the target audience of such activities.

### 3.3. Perceptions of publishing practices among STEMM ECR mentors and mentees

Publication is an important activity for the career development of ECR mentees due to the use of traditional metrics for hiring, promotion, tenure, awards, and grants [6, 11]. While the use of these traditional metrics is widely practiced, it is not well known how the relationship between ECR mentees and mentors affects ECR mentee publishing practices or perceptions of career development. Generally, it is understood that the PI or mentor directs the research objectives [32], thus some factors that affect ECR publishing practices may be influenced by individuals other than the ECR mentee. To uncover more about publishing practices among ECR mentees and mentors, a survey was conducted that investigated perceptions surrounding the types of publication important for ECR career trajectory, journal selection practices, publication practices, and the role of the mentor-mentee relationship in ECR career development. Twitter polls were also conducted to gain further insight into academic perceptions regarding publication practices by ECR mentees and mentors among the wider academic community. Furthermore, a subset of the ECR mentee and mentor survey respondents were interviewed to gain a deeper insight into the ECR mentor-mentee relationship and how these factors affect publication practices and perceptions.

#### 3.3.1. Types of publications valued by STEMM ECR mentees and mentors

While the number of manuscripts an ECR publishes is considered an important career metric [6] of which respondents to this survey were very aware (Table S6), the type of publications that are important for ECR career trajectory appear to be less well discussed in the literature. Both ECR mentees and mentors were asked about what publication types they perceive to be most important for ECR career trajectory. The results of our survey demonstrate that the overwhelming majority of ECR mentees (95.1%) and mentors (89.9%) agree that peer reviewed research papers are the most important for ECR career trajectory (Table 2). Mentees and mentors also indicate that peer reviewed narrative reviews and systematic review papers are somewhat important for career trajectory. Book chapters were also seen as somewhat important by both mentees (42.2%) and mentors (51.9%), but a greater proportion of mentees deemed book chapters not important (32.3% vs 25.9%). Generally, mentees and mentors were unsure or considered books, book chapters, case studies, technical reports, and patents less important that peer reviewed outputs. These findings appear to follow a trend reported in a large 2015 survey of faculty in the United States, which reports that researchers believe that more recognition should be awarded to traditional research publications such as journal articles, compared to other output such as data, media, or blog posts [50]. Although the respondents of that faculty survey did consider that publishing books should recognized.

**Table 2:** In the survey, ECR mentees (A; n = 106) and mentors (B; n = 28) were asked to rank these types of publication in order of importance for the career trajectory of the ECRs they mentor. Results are expressed as a percent of the total responses provided by either the mentee or mentor respondents.

|  | (A) Mentee |  |  |  | (B) Mentor |  |  |  |
| --- | --- | --- | --- | --- | --- | --- | --- | --- |
| Publication types | Not important | Unsure | Somewhat important | Very Important | Not important | Unsure | Somewhat important | Very important |
| Peer reviewed research papers | 0.0 | 2.0 | 2.9 | 95.1 | 0.0 | 7.4 | 3.7 | 88.9 |
| Peer reviewed narrative review papers | 10.8 | 22.6 | 52.9 | 13.7 | 7.4 | 3.7 | 66.7 | 22.2 |
| Peer reviewed systematic review papers (with/without meta-analyses) | 6.9 | 24.5 | 38.2 | 28.4 | 3.7 | 11.1 | 44.4 | 40.7 |
| Books | 35.3 | 13.7 | 32.3 | 18.6 | 37.0 | 29.6 | 22.2 | 11.1 |
| Book chapters | 32.3 | 9.8 | 42.2 | 15.7 | 25.9 | 14.8 | 51.9 | 7.4 |
| Case studies | 35.3 | 26.5 | 27.4 | 10.8 | 33.3 | 29.6 | 33.3 | 3.7 |
| Technical reports | 23.5 | 28.4 | 38.2 | 9.8 | 40.7 | 11.1 | 40.7 | 7.4 |
| Patents | 24.5 | 17.7 | 31.4 | 26.5 | 29.6 | 18.5 | 37.0 | 14.8 |
| Conference Abstracts | 11.8 | 11.8 | 50.0 | 26.5 | 18.5 | 3.7 | 44.4 | 33.3 |
| Conference Posters | 14.7 | 12.8 | 52.9 | 19.6 | 18.5 | 7.4 | 44.4 | 29.6 |
| Public outreach (e.g., magazine pieces, blogs, etc.) | 10.8 | 20.6 | 53.9 | 14.7 | 7.4 | 22.2 | 59.3 | 11.1 |

In this survey (Table 2), both ECR mentees (58.8%) and mentors (60%) indicated that public outreach (magazines, blogs, etc.,) were somewhat important for career trajectory, despite public outreach not being captured in the so-called traditional metrics sought by hiring or promotion committees in academia [6, 11, 51]. It is noteworthy that this was the only option presented to respondents that is perceived to be an activity by many that should be done in one’s own time and not as part of “real” research [52, 53]. In the past, outreach activities were not ordinarily incentivized or rewarded for by hiring, promotion, or tenure committees [51], despite the clear importance outreach activities are to the development of trust and understanding of science by the general public [54]. However, it is true that ECRs may be required to communicate research and the impact of their work to stakeholders such as industry partners, government agencies, and/or societies depending on their funding agencies etc., due to an increased reliance on funding from non-academic sources including the aforementioned stakeholders [55, 56].

#### 3.3.2. Journal selection practices: Prestige, readership, impact factor, and open access publishing

There is limited published information regarding how the ECR mentee-mentor relationship affects journal selection practices in the sciences. Therefore, ECR mentees and mentors were also asked to respond to statements regarding journal selection. A majority of mentees indicated that they either somewhat (30.6%) or strongly agree (57.4%) that a journal’s prestige influences their decision to submit research to a journal, which is mostly in agreement with mentors where 52% somewhat agreed and 40% strongly agreed with that statement (Table 3). Furthermore, both mentees and mentors either somewhat agreed or strongly agreed that a journal’s readership, rank, and impact factor influenced their journal sections (Table 3). Indeed, interview respondents largely indicated that high impact factor was one of the many factors they considered when publishing their manuscripts (Supplementary Table S9). These findings are in accordance with previous studies that have shown that researchers tend to aim for journals associated with prestige and high impact factors, particularly early in one’s career in order to quickly garner a reputation and improve their publication records in the highly competitive academic job market [6, 7, 57]. However, in the Twitter polls, high citation counts (41%) were considered more important for hiring academics than impact factor (13.3%), the number of publications (11.4%) or a varied skill set (34.3%). While there is some evidence that higher citation counts may be associated with publication of manuscripts in high impact factor journals [58], this ‘journal effect’ hypothesis may not always be true for all cases because there tends to be considerable variation in citation rates, especially for papers published in high-impact journals as demonstrated by Leimu and Koricheva [59]. However, differing disciplines within STEMM may experience different outcomes.

**Table 3:** ECR mentees and mentors (n = 25) were asked to provide their opinion on statements relating to publishing practices regarding journal selection. The results are expressed as a percent of the total responses provided by either the mentee or mentor respondents.

| Statements | (A) Mentee |  |  |  |  | (B) Mentor |  |  |  |  |
| --- | --- | --- | --- | --- | --- | --- | --- | --- | --- | --- |
|  | Strongly disagree | Somewhat disagree | Neither agree nor disagree | Somewhat agree | Strongly agree | Strongly disagree | Somewhat disagree | Neither agree nor disagree | Somewhat agree | Strongly agree |
| A journal's prestige influences the decision to submit the research | 0.0 | 1.0 | 11.2 | 30.6 | 57.4 | 4 | 0 | 4 | 52 | 40 |
| A journal's readership influences the decision to submit the research | 0.0 | 2.0 | 16.3 | 32.6 | 48.9 | 0 | 4 | 8 | 34 | 54 |
| Rank and impact factor influences the decision to submit the research | 1.0 | 3.1 | 11.2 | 34.6 | 50.0 | 40 | 0.0 | 4 | 48 | 44 |
| A journals accessibility (open access or not) to the reader influences the decision to submit the research | 1.0 | 8.2 | 21.4 | 30.6 | 38.8 | 0.0 | 8 | 12 | 56 | 24 |
| The cost of article processing fees determines where the research is submitted | 8.2 | 12.2 | 25.5 | 29.6 | 24.5 | 0.0 | 8 | 12 | 32 | 48 |

Mentees and mentors were also asked to write about if they had journals or publishers that they favored and to say why (Figure 3). Word cloud analyses show that both mentors and mentees placed an emphasis on “impact”. Of the 79 mentees that responded (Figure 3A), “impact” was mentioned 18 times along with publishers such as Nature (15 times), Science (10 times), and Elsevier (4 times). Mentors mentioned “impact” 6 times out of 20 respondents. As shown in Figure 3, respectability and impact factor were also recurring words. One mentee wrote that they favored “those with higher impact factor … those with positive reputation in terms of research conduct…”, which was echoed by another mentor who said they prefer “high impact [journals]…due to their reputation among funders and colleagues” (Table S2).

**Figure 3:**
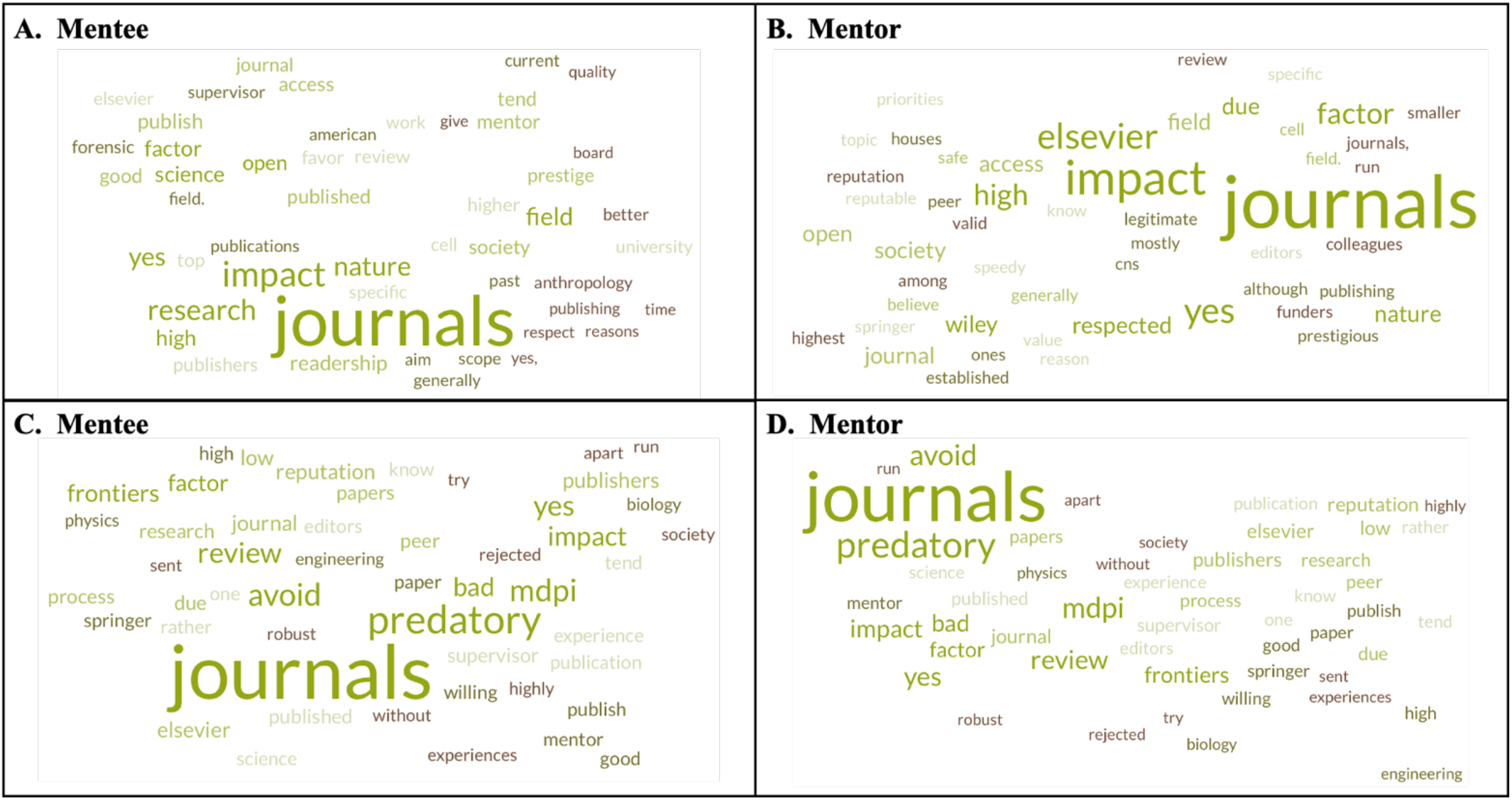
Word clouds were generated from the top 50 most frequently used words by mentees (n = 79 respondents) or top 25 most frequently used words by mentors (n = 20 respondents) in response to questions regarding journal selection. Mentees were asked: (A) Are there research journals or publishers that you or your (supervisor) tend to favor? If so, why? Mentors were asked: (B) Are there research journals or publishers that you tend to favor? Why/why not? Mentees were also asked: (C) Are there research journals or publishers that you or your (supervisor) tend to avoid? If so, why? Mentors were asked: (D) Are there research journals or publishers that you tend to avoid? Why/why not? The word cloud were generated using an online generator [77]. The darker the intensity of the green colors and larger the words, the greater the frequency the word was used by respondents.

Indeed, mentees seemed to strongly agree that prestige influenced their journal sections (57.4%), whereas the majority of mentors only somewhat agreed (52%) (Table 2). Society journals also received a notable mention by both mentees and mentors as journals that are favored. One mentor stated that they “give back to the community” (Table S2). Additional data regarding mentor mentee responses is available in the supplementary materials (Tables S2 and S4). These findings are in accordance with previous research that found that the reputation of a journal was an important factor influencing researchers’ journal selection [60].

The scope of the journal is also highlighted in the literature as an important factor that influences journal selection decisions by researchers [60]. Indeed, when writing about journal preferences in the survey, one mentee highlighted that they prefer to select journals for their “target audience” and another mentioned that they tend to avoid “journals that do not fit their scope” (Supplementary Tables S1 and S3). Other factors that have previously been identified as important for journal selection decisions include reliability of reviewing and usefulness of reviewers’ feedback [60]; however, these factors did not feature heavily in the responses to the survey or the interviews.

On the other hand, the Twitter polls indicated that readership (44.7%) and accessibility/open access (32.9%) were considered more influential than impact factor (5.9%), and article processing charges (16.5%) when choosing a journal (total of 85 votes; Table 1). In the survey, ECR mentees (38.8%) strongly agreed that a journal’s accessibility, whether it is open access or not, is an important factor for journal selection, whereas a majority of mentors (56%) deemed accessibility as somewhat important (Table 3).

Previous research has indicated that younger researchers prefer open access publishing versus more established academics [57] as evidenced by the 64% of ECR faculty members in a large survey in the United states who strongly agreed that they would be happy to see traditional journal subscription models replaced by open access models entirely [50]. Despite research showing that open access publishing as a principal is widely endorsed, in practice it was deemed one of the least important factors that influenced author journal selection [61–63]. One potential reason may be because of the high article processing charges (APC) associated with publishing in open access journals, when some journals offer a subscription model that does not cost the authors to publish [64]. However, Nicholas, Rodríguez-Bravo [6] postulates that ECRs are preoccupied with chasing higher impact factor journals that have traditionally not been open access in practice. In recent years, publishers considered prestigious such as Nature Publishing Group have opted to offer both the subscription model and the open access models for most journals, with some journals deemed fully open access, which requires authors to pay APC to be published [65]. In the survey, while over half of mentees somewhat agreed (29.6%) or strongly agreed (24.5%) that the cost of APC determines journal selection, a greater proportion of mentors indicated that they either somewhat (32%) or strongly agree (48%).

Both ECR mentors and mentees were also asked in the survey if there was journals or publishers that they tended to avoid. One mentee wrote “we tend to avoid journals with high article processing charges”, which was echoed by 6 ECR mentors who said they tend to avoid journals with “high publishing fees” with one mentioning it was due to “no funds for that” (Supplementary Table S4). Mentee interviewee 1 also indicated that they preferred journals that did not charge APC due to lack of funding, but that they had difficulty matching their submissions to the scope of journals due to their limited options (Supplementary Table 17).

ECR mentors and mentees also specified several open access publishers in response to a question regarding whether there were journals that they tended to avoid (Figure 3C and 3D, Table S2 and S4). Respondents expressed dissatisfaction with journals that used “pay to publish” models and had “poor reputations” as indicated by the word clouds (Figure 3, Tables S3 and S4). These responses were mostly associated with who respondents perceived to be “predatory publishers”. As summarized by Gallent Torres [66], predatory journals are generally defined by authors as journals that exploit the open access model unprofessionally to achieve economic gains at the expense of the quality standards of scientific publications. Indeed, predatory publishing practices were mentioned by 10 ECR mentors and 18 ECR mentees (Figure 3). In recent years there has been growing concern and criticism of some open access publishers mentioned by the respondents (Figure 3, Tables S2 and S4), likely due to their exponential growth [67, 68] and the rise of mega-journals that publish vast quantities of manuscripts [69, 70]. However, Nicholas, Rodríguez-Bravo [6] has reported that some ECR mentees in some fields misunderstand and conflate open access publishing with predatory publishing. Additionally, some respondents of the survey in this study who had expressed concerns regarding specific publishers also admitted that they have previously published in some of these journals (Supplementary Tables S3 and S4). This may not be surprising as many of the publishers mentioned by the respondents are now among the leading growing publishing groups of scientific literature, challenging the more established and larger publishers like Elsevier [71]. These include publishers such as MDPI and Frontiers, who are the subject of intense debate due to their exponential growth, prolific use of special issues, and concerns about peer review and editorial rigor [72–76]. This raises numerous questions regarding whether publishing in so-called predatory journals or even mega-journals may affect future ECR career outcomes, which were not addressed in the current study. Indeed, there has been recent discussion whether MDPI and Frontiers journals may be downgraded to the lowest level (level 0) in the Finnish Publication Forum, which is a quality of research assessment for hiring and promotion [76].

#### 3.3.3 Manuscript handling and authorship practices among ECR mentees and mentors

Manuscript preparation can be a daunting task for ECR mentees [78]. While there are countless manuscripts available that provide ECRs and more senior researchers with guidance for writing scientific manuscripts [78–80], research regarding the responsibilities of ECR mentees and mentors in these processes is largely lacking from the literature. To gain further insight about how ECR mentees and mentors approach manuscript preparation and authorship practices, a series of questions were devised in the surveys and interviews.

As shown in Table 4, ECR mentors and mentees largely agree on who is responsible for certain tasks with regards to manuscript preparation. For example, over 88% of ECR mentees indicated that they somewhat or strongly agree that they lead the writing of the first draft of a manuscript, and mentors confirm their responses (86.9%). Likewise, ECR mentees and mentors mostly agree that mentors contribute to the drafting of manuscripts. However, ECR mentees (66.5%) somewhat or strongly agree that their mentors are reliant on them to write these manuscripts. Moreover, half of the mentees (51.1%) either strongly or somewhat disagreed that their mentor contributes to the formation of tables or figures in a manuscript, which differs from mentors (69.6%) who believe that they do contribute to these scholarly activities.

**Table 4:** ECR mentees (A; n = 94) and mentors (B; n = 23) were asked to respond to similarly phrased statements regarding manuscript preparation. The results are expressed as a percent of the total responses provided by either the mentee or mentor respondents.

|  | Percent of respondents (%) |  |  |  |  |
| --- | --- | --- | --- | --- | --- |
| **(A) Mentee statements** | **Strongly disagree** | **Somewhat disagree** | **Neither agree nor disagree** | **Somewhat agree** | **Strongly agree** |
| I lead the writing of the first draft of a manuscript | 6.4 | 1.1 | 4.3 | 12.8 | 75.5 |
| My mentor contributes to the drafting of manuscripts | 3.2 | 10.6 | 13.8 | 30.9 | 41.2 |
| My mentor is receptive to critique and feedback to sections of a manuscript written by my mentor | 1.0 | 5.4 | 22.6 | 36.6 | 34.4 |
| My mentor is reliant on me to write manuscripts for publication | 6.4 | 6.4 | 20.4 | 21.25 | 45.2 |
| I am responsible for reviewing manuscripts prior to submission | 2.1 | 5.3 | 9.6 | 30.9 | 52.1 |
| My mentor contributes to the creation of figures and tables | 30.9 | 20.2 | 15.9 | 19.1 | 13.8 |
| My mentor's feedback is clear and easy to follow | 4.3 | 8.5 | 8.5 | 31.9 | 46.8 |
| **(B) Mentor statements** | Percent of respondents (%) |  |  |  |  |
|  | **Strongly disagree** | **Somewhat disagree** | **Neither agree nor disagree** | **Somewhat agree** | **Strongly agree** |
| My ECRs lead the writing of the first draft of a manuscript | 8.7 | 4.4 | 0.0 | 30.4 | 56.5 |
| I contribute to the writing of the first draft of a manuscript with my ECRs | 13.0 | 8.7 | 4.4 | 26.1 | 47.8 |
| My ECRs willfully provide critique and feedback on my writing sections | 0.0 | 21.7 | 8.7 | 30.4 | 39.1 |
| I review the manuscripts prior to submission with consultation of my ECRs | 0.0 | 0.0 | 4.4 | 4.4 | 91.3 |
| I contribute to the creation of tables and figures | 13.0 | 4.4 | 13.0 | 26.1 | 43.5 |
| I try to give clear feedback to my ECRs | 4.4 | 0.0 | 0.0 | 21.7 | 73.9 |

Questions were also devised in the survey and interviews to determine if there were varying perceptions regarding authorship practices among ECR mentees and mentors. Authorship is an important output for scholars, but particularly for ECRs who are reliant on publications for future job prospects [9].

In the survey, mentees (78.5.1%) and mentors (96%) somewhat or strongly agreed that authorship is dealt with fairly and equitably. However, a minority of ECRs have somewhat or strongly indicated their involvement in authorship disputes with either their mentor (13.2%) or with someone else (30%). However, only one mentor indicated that they somewhat agreed that they had authorship disputes with their mentees and only 12% of mentors somewhat agree that members of their team have had disagreements with each other regarding authorship. In contrast, a poll conducted on Twitter indicated that almost half (47.5%) of the respondents (n = 80) specified that they have had authorship disputes. Indeed, some of the ECR mentee and mentor interviewees reported that they had either witnessed or been involved in authorship disputes in the past (Supplementary Table S9). Mentee interviewee 2 had indicated that on more than one occasion they had issues with authorship. In particular, they discussed how their mentor expressed “favoritism” for other mentees, which led to authorship disputes. Mentee interviewee 2 described the ordeal as “exhausting” and said “you don’t think it matters, but it does. It does matter when you’re applying for things in the future” (Supplementary Table S9). From another perspective, mentor interviewee 1 discussed the “competition” that exists regarding authorship between their ECRs and said that there has been disputes in the group, but they try to minimize the “friction as early as possible” by agreeing on authorship early to “keep a balance” regarding who is first or second authors on manuscript (Supplementary Table S9, mentor 1). Mentor interviewee 2 also expressed that they had authorship dispute “multiple times” in their past. They indicated that they had been excluded as an author on papers that they had contributed to (Supplementary Table S9, mentor 2). This is not uncommon, and these individuals are sometimes referred to as “ghost writers”, individuals who contributed to the research but were excluded from the final manuscript. In one study, as many as 8% of corresponding authors in leading medical journals admitted to removing individuals from authorship [81]. Table 5 shows that 15.3% and 17.4% of ECR respondents strongly agree that they were unfairly removed from the authorship of a manuscript or witnessed research misconduct respectively. Smith et al., [82] surveyed 6,700 international researchers in the United States and found that 46.6% of researchers had experienced disagreements about author naming and 37.9% had disputes about the order of authors on manuscripts [82]. These findings appear to be closer to the findings of our Twitter polls (Table 1).

**Table 5:** ECR mentees (A; n = 98) and mentors (B; n = 25) were asked to respond to similarly phrased statements regarding publication ethics and authorship practices. The results are expressed as a percent of the total responses provided by either the mentee or mentor respondents.

|  | Percent of respondents (%) |  |  |  |  |
| --- | --- | --- | --- | --- | --- |
| **(A) Mentee statements** | **Strongly disagree** | **Somewhat disagree** | **Neither agree nor disagree** | **Somewhat agree** | **Strongly agree** |
| I am aware of the CRediT author statement or similar author taxonomies | 21.4 | 7.1 | 12.2 | 19.4 | 39.8 |
| My mentor is fair and equitable regarding authorship | 2.0 | 6.1 | 13.3 | 22.4 | 56.1 |
| I have had disputes regarding authorship with my mentor | 59.2 | 13.3 | 14.3 | 6.1 | 7.1 |
| I have had disputes regarding authorship with individuals other than my mentor | 42.9 | 12.2 | 15.3 | 16.3 | 13.7 |
| I have witnessed research misconduct | 46.9 | 13.3 | 6.1 | 15.3 | 17.4 |
| I have shared or would share first authorship with another researcher | 7.1 | 1.0 | 12.2 | 26.5 | 53.1 |
| I have been asked/forced to change my authorship position on a manuscript | 52.0 | 13.3 | 9.2 | 13.3 | 12.2 |
| I was unfairly excluded from a manuscript(s) authorship | 57.1 | 13.3 | 6.1 | 8.2 | 15.3 |
| The number of authors on a manuscript matters | 24.5 | 13.3 | 23.5 | 26.5 | 12.2 |
| My mentor has discussed authorship with me prior to the initiation of a project | 15.3 | 14.3 | 17.4 | 23.5 | 29.6 |
|  | Percent of respondents (%) |  |  |  |  |
| **(B) Mentor Statements** | **Strongly disagree** | **Somewhat disagree** | **Neither agree nor disagree** | **Somewhat agree** | **Strongly agree** |
| I have discussed the CRediT author statement or similar contributor roles taxonomies with my ECRs | 12 | 4 | 12 | 28 | 44 |
| I am a fair and equitable mentor when deciding who is an author on a manuscript | 0.0 | 4 | 0.0 | 24 | 72 |
| I have had disputes or disagreements regarding authorship with my ECRs | 68 | 24 | 4 | 4 | 0.0 |
| Members of my team have had disagreements with each other regarding authorship | 64 | 12 | 12 | 12 | 0.0 |
| I have witnessed research misconduct | 40 | 24 | 12 | 12 | 12 |
| I have excluded ECRs from a manuscript(s) authorship because I thought they did not contribute enough | 60 | 16 | 16 | 4 | 4 |
| It is important to limit the number of authors on my manuscripts | 56 | 8 | 24 | 12 | 0.0 |
| I discuss authorship with my ECRs prior to the initiation of a project or manuscript with them | 4 | 4 | 4 | 32 | 56 |
| I disclose all conflicts of interest accurately in a manuscript | 0 | 0 | 0 | 12 | 88 |

Collectively, it would appear that authorship disputes are somewhat more common than our survey data suggests. However, over a quarter of ECR survey respondents indicated that they were not aware of the CRediT author statement or similar contributor roles taxonomies used by most publishers (Table 5) and only 72% of mentors indicated that they somewhat oe strongly agreed that they have discussed the CRediT authorship statement with their ECRs. These statements are a means to present an accurate and detailed representation of individual author contributions of a published work. This is a rather worrying finding as almost all reputable journals require corresponding authors (mentors) to acknowledge that the CrediT author statement was followed. One must question if there was greater knowledge, understanding, or enforcement of these contributor role taxonomies, would there be less prevalence of authorship disputes?

### 3.4. Communication is a key factor in the relationship between ECR mentees and mentors that influences ECR publishing practices

As discussed previously, ECR mentees tend to rely on mentorship, supervision, and training facilitated by a more experienced researcher, usually a PI, who directs the objectives of a research group, which may affect the publishing practices and therefore the career development of the ECRs mentees [32, 83] and mentors [35]. To further understand what factors or processes are required to maintain these relationships, survey participants were asked to provide information regarding what they believe is important regarding a healthy ECR mentor-mentee relationship.

Generally, the mentee survey respondents and interviewees placed a significant emphasis on communication as an important characteristic of a good mentor who may assist in advancing their careers, which is in accordance with previously published literature [84, 85]. This was evident in the word cloud analysis in Figure 4A, where communication was mentioned by 29 of the 68 mentee respondents with regards to what characteristics mentee’s thought are necessary for a healthy ECR mentor-mentee relationship (Table 6). Mentors tend to agree with mentees, as 7 out of 18 mentor survey respondents noted that communication is required for a healthy ECR mentee-mentor relationship (Table 6B and Figure 4B).

**Figure 4:**
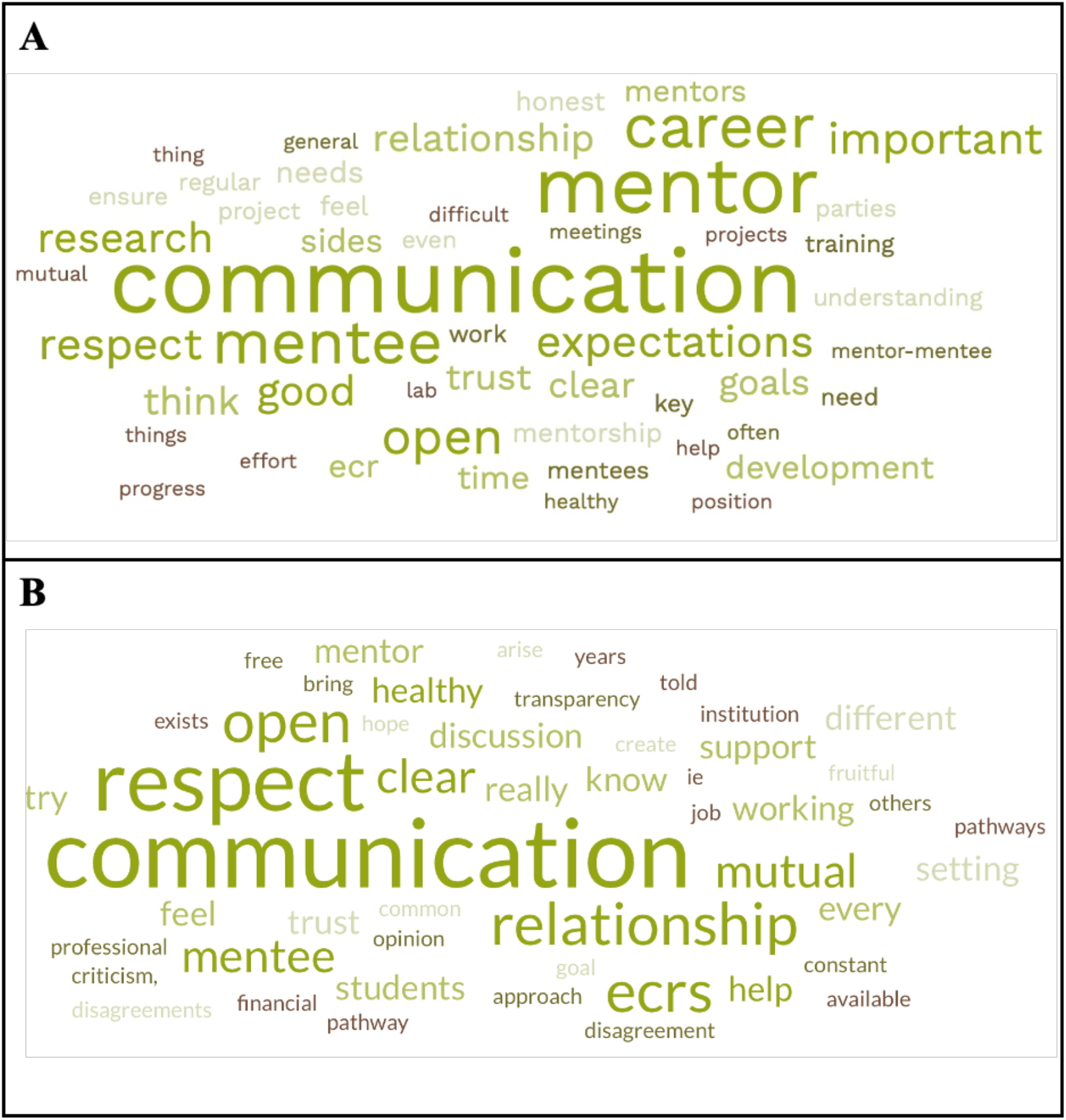
Word clouds were generated from the top 50 most frequently used words in response to the following task assigned to both mentees (A; n = 68 respondents) and mentors (B; n = 18 respondents) in the survey: In the space provided, write a few lines about what you believe is important regarding a healthy ECR mentor-mentee relationship. The word cloud were generated using an online generator [77]. The darker the intensity of the green colors and larger the words, the greater the frequency the word was used by respondents.

**Table 6:**
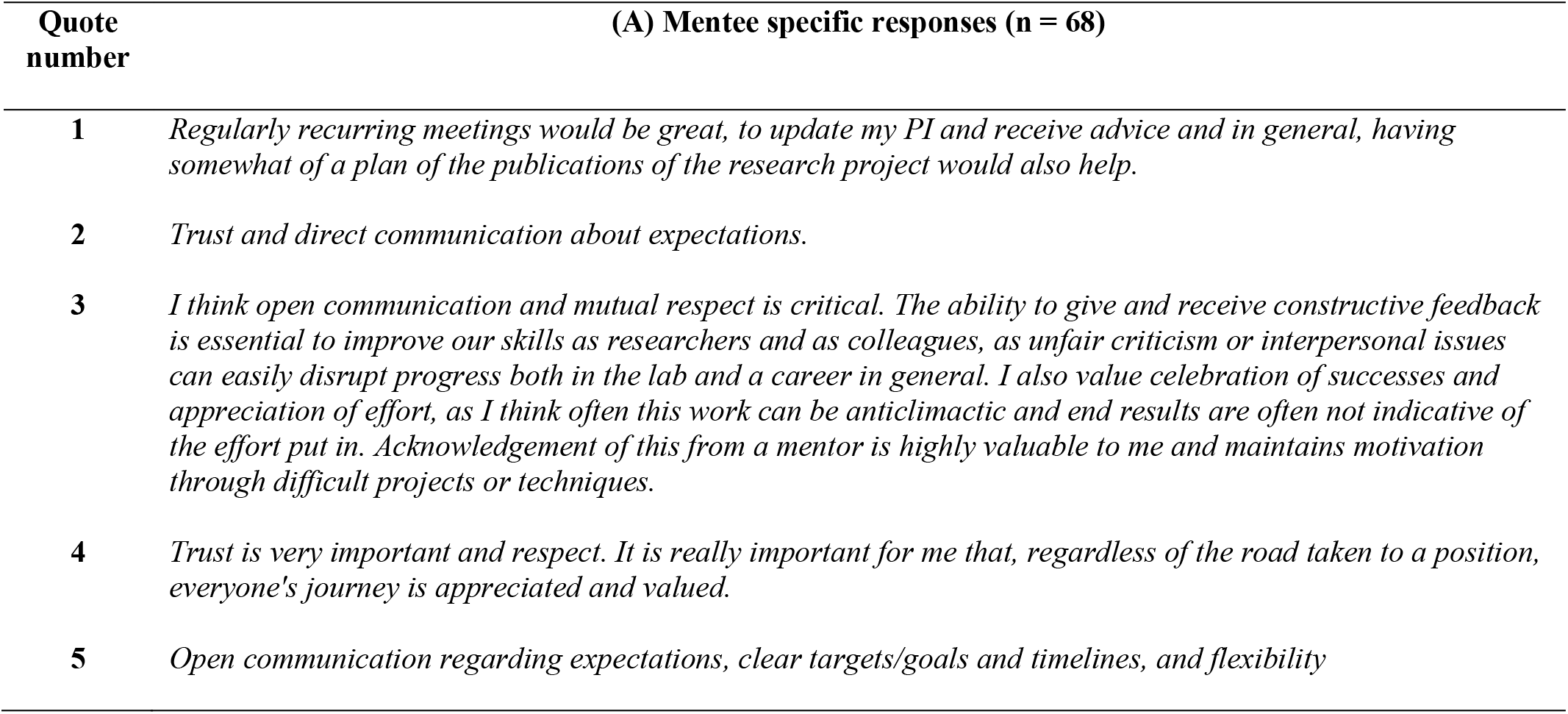
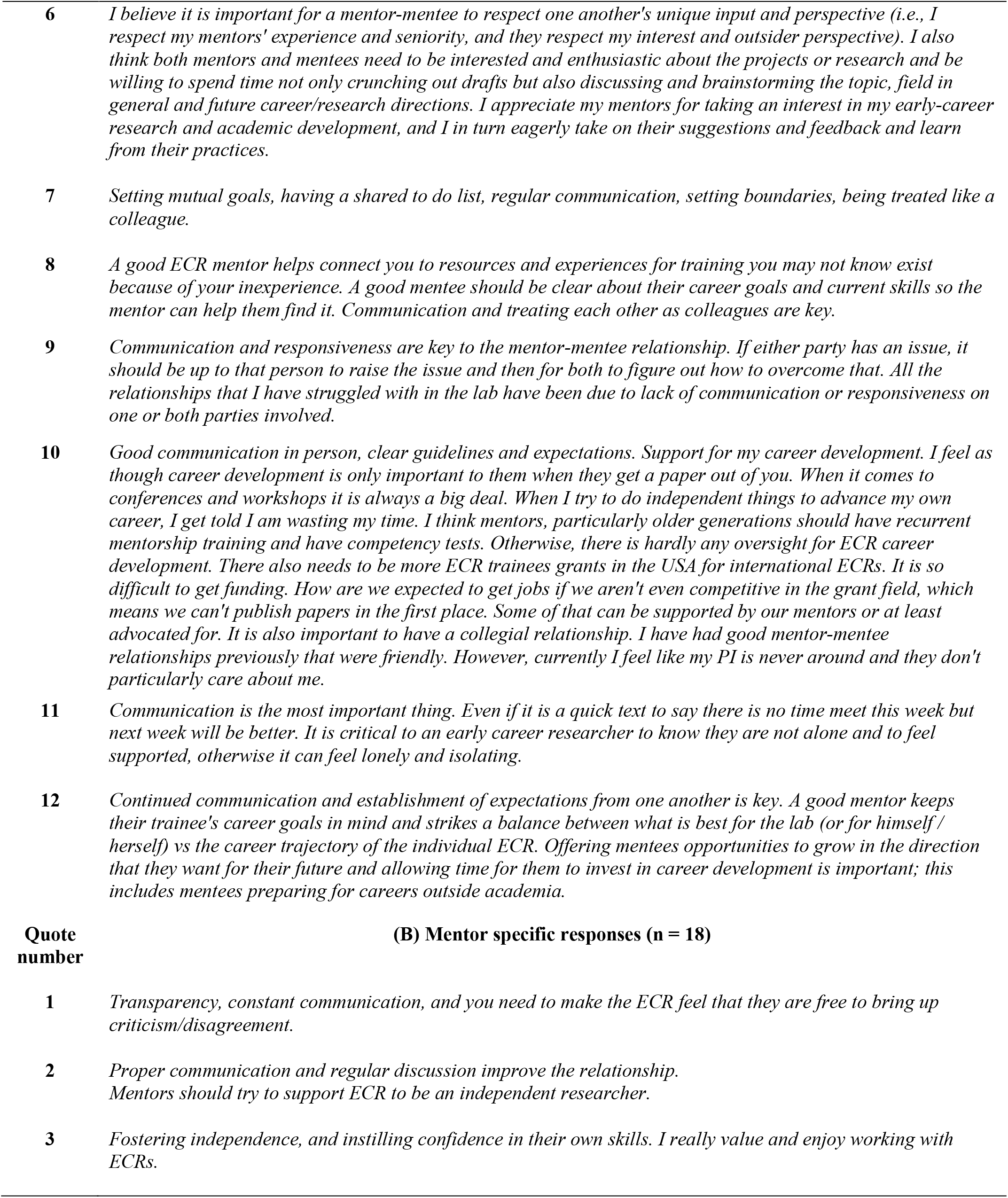
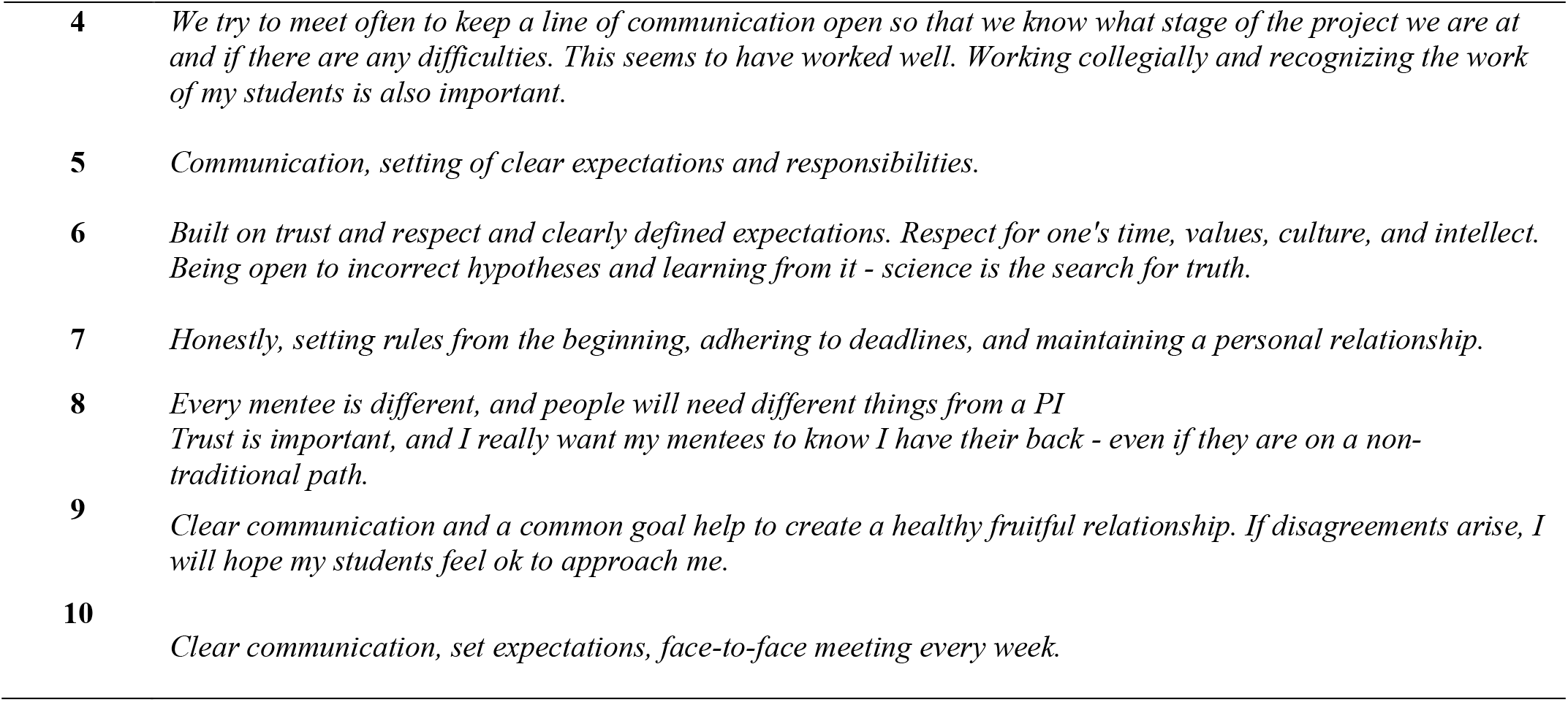
ECR mentees and mentors were asked to write a few lines about what they believe is important regarding a healthy ECR mentor-mentee relationship in the survey. Below are some of the typical responses received from mentees (A) and mentors (B) who responded.

Mentees also highlighted that “regular communication”, “trust’, “mutual, respect”, and “treating each other as colleagues” were important features of a healthy mentee-mentor relationship (Table 6A and Figure 4A). These views were also largely shared by the mentors as evidenced by the broad overlap of the frequency of words identified in the word cloud analyses (Figure 4). Additionally, in the survey, mentors highlighted that it was important to work “collegially”, maintain a “personal relationship” with mentees, and be open to discuss “criticism/disagreements” (Table 6B and Figure 4B). These findings are corroborated by the data showing that 87% of mentee survey respondents (n = 100) indicated that they somewhat or strongly agreed that they had a collegial relationship with their mentors (Supplementary table S5).

Planning and communicating clear expectations also emerged as an important factor that both mentees and mentors’ thought were important for career development. One mentee indicated that “regularly recurring meetings would be great, to update my PI and receive advice and in general, having somewhat of a plan of the publications of the research project would also help” (Table 6). Research has shown that communicating expectations around writing and publication strategies or conducting individualized development plans (IDPs) between mentees and mentors may affect ECR publication outputs and potentially career progress [86]. Indeed, mentee respondents of this survey reported that they have some form a career development plan (63.8%), which is a practice that mentors somewhat or strongly encourage (69.1%; Supplementary table S6). Research regarding IDPs and career development plans acknowledge that communication between mentees and mentors is necessary to formulate and execute these plans, and the IDPs themselves can act as a form of communication between mentees and mentors to allow for dialogue regarding shared expectations and aspirations [87]. These type of career development activities can also be further enhanced by formal mentoring programs for mentors to upskill [88]. However, it appears that these mentor training opportunities are not widespread, as mentor respondents of the survey said that “there is no training to be a mentor” (Supplementary Tables S7 and S8) and an interviewee indicated that they had asked their “department to organize workshops or even courses on how to supervise students” (Mentor interviewee 2; Supplementary Table S10B).

Communication between ECR mentees and mentors can reciprocally benefit the career development of both parties in several ways. As indicated by mentor interviewee 1 regarding mentoring ECRs, “you build a personal relationship with this person by trying to help him or her with his or her career” (Table S10B). However, communication also benefits the mentors who gain from “comments exchanged during discussion” (Mentor interviewee 1; Table S10B). Respectful communication and collegiality between ECR mentees and mentors also affect career development through the process or writing and preparing research for publication. Collegiality can allow for meaningful feedback between mentees and mentors when preparing manuscripts, which is an important part of writing scientific manuscripts and is an opportunity for career development. Mentees can exchange dialogue with their mentors to improve their writing and data interpretation skills along with improving the manuscript for submission on the journey [80]. In the survey, the majority of ECR mentees (71%) reveal that their mentors are receptive to feedback during the writing process (Table 4). Research shows that ECRs value feedback from their mentors and that relationships developed from these communications are important for ECR productivity [34] as publications play a significant role in ECR career development [89].

Considering the consensus among mentors and mentees regarding the importance of regular communication in Figure 4, there are some disparities in the reporting of the frequency of face-to-face communication and online communication between mentee and mentor respondents. While two-thirds of mentors indicate that they have face-to-face communication either once or more than once per week, the majority of mentees indicate that they have face-to-face communication daily (4%), once per week (25%), more than once per week (15%), with a fifth of mentees indicating that they did not often have face-to-face communication (Figure 5). Similarly, the majority of mentors indicated that they have online communication with their mentees either daily (37%) or more than once per week (29.6). However, only 16% of mentees indicated that they had daily contact with their mentors, and that it was more likely to be once per week (21%) or more than once per week (24%). Table 7 and Table 8 contain quotes from mentees and mentors respectively regarding communication in the mentee-mentor relationship. It is clear that different mentees and mentors have varied experiences regarding communication.

**Figure 5:**
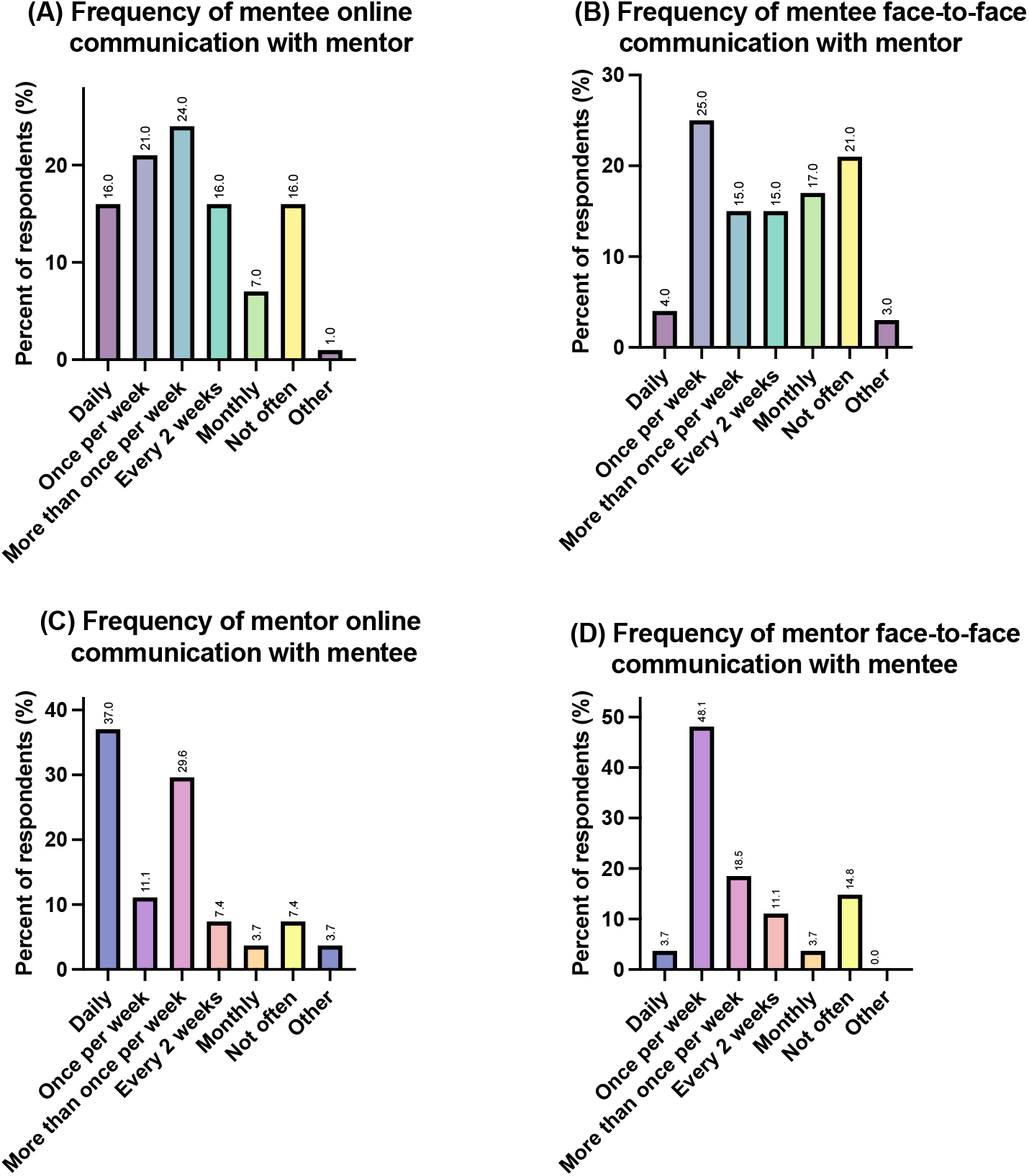
Mentees’ and mentors’ perception of their frequency of contact online (panels A and C; n = 100 respondents) and face-to-face (panels B and D; n = 27 respondents).

**Table 7:** Mentees were asked “overall, do you think you and your mentor communicate well? Are there any positive or negative points that you would like to share?” These exemplary responses were selected from a total of 76 responses recorded.

| Quote number | Specific responses |
| --- | --- |
| 1 | *I think we have good communication and are accessible for each other. Our lab uses slack as a direct messaging system, which is very helpful for short messages or quick checks, and we also have dedicated half hour time slots each week to meet 1-on-1. We don't always fill that time, but it is very helpful to know that there is time allocated for discussion about any aspect of my work or career progression. I also have annual reviews which are more detailed reflections on progress e.g., publications, grant writing, presentations.* |
| 2 | *I think my mentor and I communicate well and on a professional yet friendly level. There is no superiority complex, which I believe is important in order to have open and honest conversations. We also understand and respect each other's boundaries.* |
| 3 | *My supervisors are excellent, particularly my primary one, but we don't meet often enough primarily because of job demands, the research suffers.* |
| 4 | *I think we could communicate better. We are actually not too different in age, but our career trajectory has been very different. I took a career break and changed field, so while I am not as advanced in this field I have a lot of experience of leading research, winning grants, and supervising students, just in a different field. I think he finds it difficult to not to treat me as very junior and finds it more difficult to accept my ideas as valid.* |
| 5 | *I feel my mentor and I communicate well. We have consistent communication that balances my need for guidance and space. I also feel comfortable talking with her; I do not dread our meetings or feel anxious, afraid, or belittled.* |
| 6 | *No, my mentor is very busy. He wants to be there for me in theory, but in practice I don't feel comfortable talking with him about problems and issues that arise in my work. He has a different approach than me and has handled conflicts very poorly.* |
| 7 | *The principal investigator of my project has very little interest (or time) towards me and the other Ph.D. students. We sometimes do not discuss issues for months and she has no overview of what I'm doing.* |
| 8 | *My mentor's leadership style is generally hands-off. He is a very busy person, so I do my best to make the most of allotted meeting times (i.e., lab meetings) and rarely request additional meetings. He is always responsive over email though, so this has been an effective means of regular communication.* |
| 9 | *I'm just a cog in the machine. He's nice and smart though* |
| 10 | *I currently have 2 mentors, one at each of my institutions. I believe i communicate with both of my mentors well. I am enjoying experiencing the different styles of communication from people of different cultures. For example, as an Irish person trained in Ireland, I have the habit of being overly nice or not getting straight to the point. This is something I might need to work on for my future career to be able to stand up for myself and communicate clearly with everyone. One of my mentors at the moment is a great straight talker, not overly harsh, but straight to the point. I was not used to this at the beginning, but now I see that in the end it saves everybody time and it is a more efficient way of working.* |
| 11 | *Yes, as the first postdoctoral researcher in my current laboratory, my mentor and I have navigated many of the challenges of establishing a research program in a cooperative fashion. In some cases, this has resulted in additional stress to publish, since my advisor will be up for tenure review soon. Overall, we have developed a strong working relationship that recognizes our mutual strengths and differences in our approaches/research interests.* |
| 12 | *No... I expected to do good research and worked hard, earned my independent fellowship but in return she stole my ideas from lab meeting and published as her own. Additionally, I faced racial comments and discrimination. I complained to the XX institute, but I got no help. Thankfully in the time of mental breakdown, the NHS provided me the treatment of depression. It took more than 3 years to overcome from that mental trauma. I am still trying to overcome this trauma as I aim to establish my own independent group in XXXXX. The lesson learned is to support other students and postdoc with kindness, better research environment, funds, equality and support and hope they (or anyone) will not suffer like I suffered in my past.* |

**Table 8:**
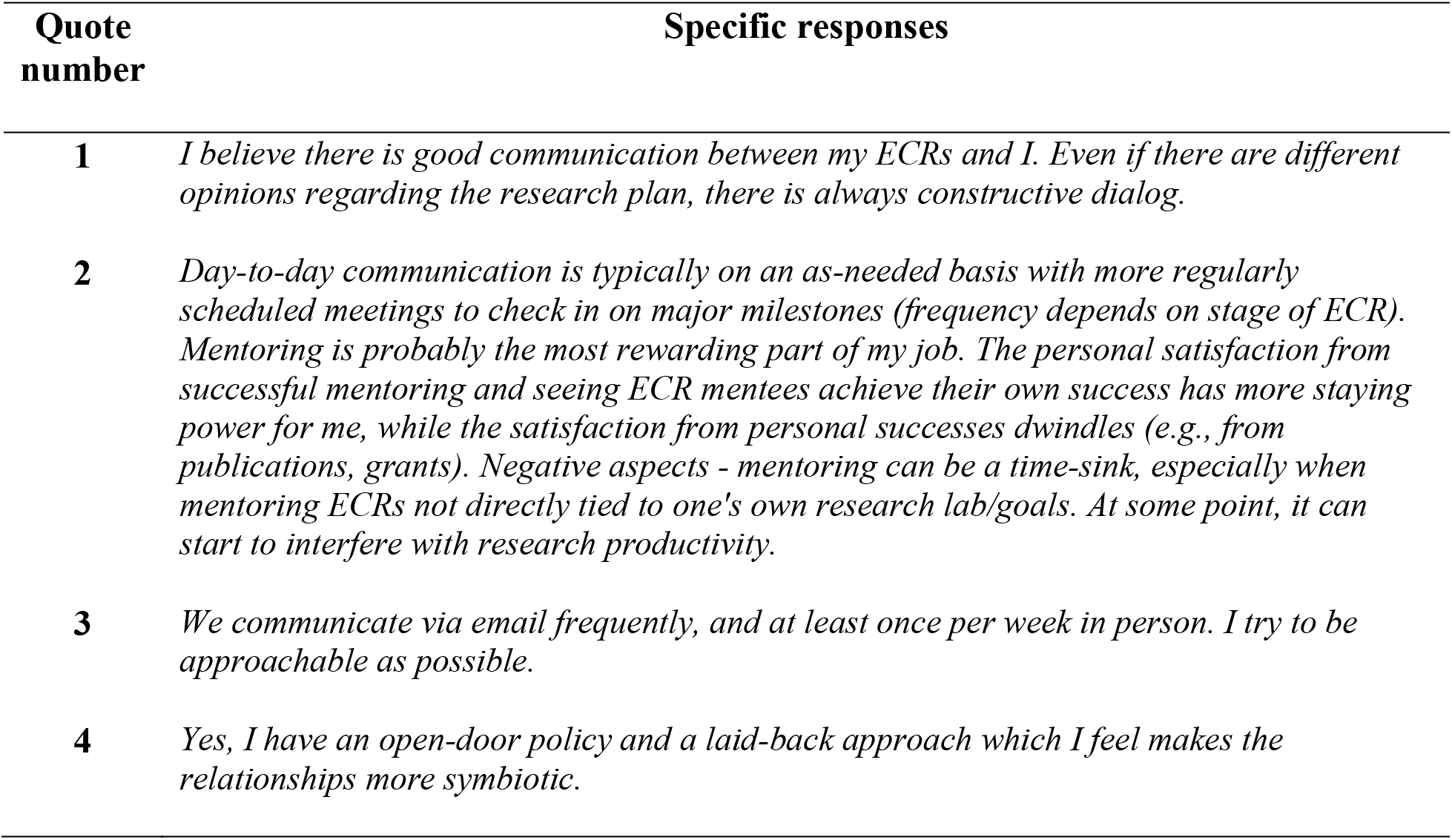
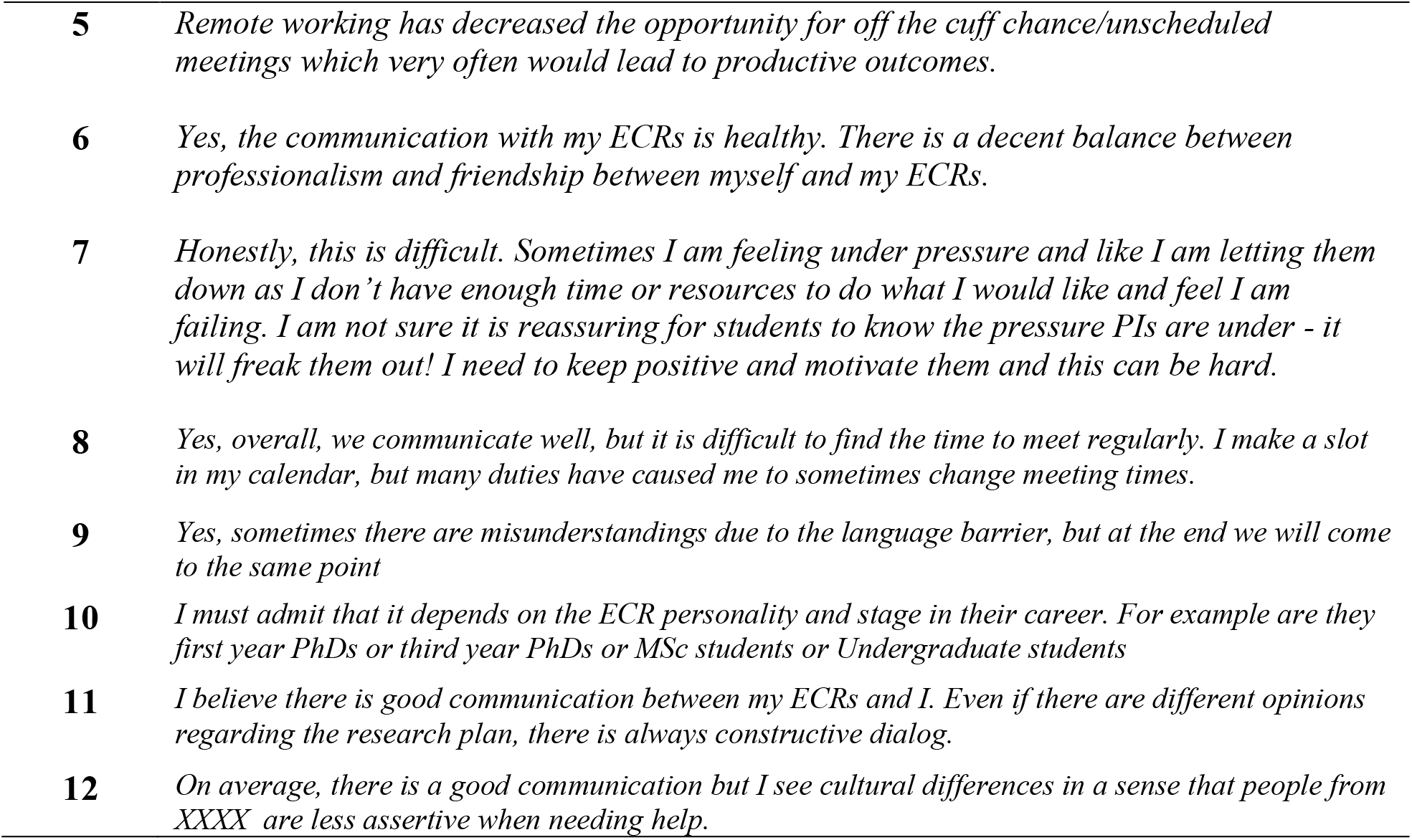
Mentors were asked: Overall, do you think you and your ECRs communicate well? Are there any positive or negative points that you would like to share? A selection of exemplary responses is presented below from a total 20 responses recorded.

| Quote number | Specific responses |
| --- | --- |
| 1 | *I believe there is good communication between my ECRs and I. Even if there are different opinions regarding the research plan, there is always constructive dialog.* |
| 2 | *Day-to-day communication is typically on an as-needed basis with more regularly scheduled meetings to check in on major milestones (frequency depends on stage of ECR). Mentoring is probably the most rewarding part of my job. The personal satisfaction from successful mentoring and seeing ECR mentees achieve their own success has more staying power for me, while the satisfaction from personal successes dwindles (e.g., from publications, grants). Negative aspects - mentoring can be a time-sink, especially when mentoring ECRs not directly tied to one's own research lab/goals. At some point, it can start to interfere with research productivity.* |
| 3 | *We communicate via email frequently, and at least once per week in person. I try to be approachable as possible.* |
| 4 | *Yes, I have an open-door policy and a laid-back approach which I feel makes the relationships more symbiotic.* |
| 5 | *Remote working has decreased the opportunity for off the cuff chance/unscheduled meetings which very often would lead to productive outcomes.* |
| 6 | *Yes, the communication with my ECRs is healthy. There is a decent balance between professionalism and friendship between myself and my ECRs.* |
| 7 | *Honestly, this is difficult. Sometimes I am feeling under pressure and like I am letting them down as I don't have enough time or resources to do what I would like and feel I am failing. I am not sure it is reassuring for students to know the pressure PIs are under - it will freak them out! I need to keep positive and motivate them and this can be hard.* |
| 8 | *Yes, overall, we communicate well, but it is difficult to find the time to meet regularly. I make a slot in my calendar, but many duties have caused me to sometimes change meeting times.* |
| 9 | *Yes, sometimes there are misunderstandings due to the language barrier, but at the end we will come to the same point* |
| 10 | *I must admit that it depends on the ECR personality and stage in their career. For example are they first year PhDs or third year PhDs or MSc students or Undergraduate students* |
| 11 | *I believe there is good communication between my ECRs and I. Even if there are different opinions regarding the research plan, there is always constructive dialog.* |
| 12 | *On average, there is a good communication but I see cultural differences in a sense that people from XXXX are less assertive when needing help.* |

## 4.0 Critical insights and recommendations for future ECR mentorship and career development

The overarching aim of this study was to determine how the ECR mentor-mentee relationship affects ECRs and their perceptions of career development in the STEMM disciplines. Two additional research questions were developed.

The first research question was who do the STEMM disciplines (ECR mentees and mentors) currently perceive to be ECRs. In this study, it was determined that there is a broad varying opinion regarding who is perceived to be an ECR by both mentees and mentors. Mentees mostly perceive Ph.D. researchers and postdoctoral research as ECRs, and while mentors agree, they also perceive new PIs (0-2 years’ experience) to be ECRs. This likely stems from the varying definitions that exist among academic institution, funding bodies, and government agencies for the purpose of hiring, promotions, and awarding grants or awards [3, 6, 7]. It is important moving forward that academia, government, and granting agencies internationally come to a consensus regarding the definition of an ECR due to the implications the definition may have on the criteria for a grant or job prospects that affect the career development of boths ECR and their mentors. Many important career development grants such as the National Institutes of Health K99/R00 Pathway to Independence Award for example [90], limit applicants to within 4 years of completing their Ph.D. [10]. Some funders, including the European Reseach Counsil Starting Grant allows ECRs to apply > 2 years but ≤ 7 post Ph.D graduation [91]. While this may be plenty of time in some disciplines, in others where longitudinal human research or animal studies may be involved, it may not be enough time to publish research or findings that may support one’s application. These inequalities may disadvantage ECRs and impact their career development in some fields of research.

The second research question addressed in this study was to determine what characteristics of the ECR mentee-mentor relationship are perceived to be important for ECR publishing practices and career progression. For better or worse, the so-called traditional metrics of publishing are interlinked with career progression of both ECR mentees and mentors, and are considered predictors of future success in academia [9, 21]. The survey and interview data obtained in this study indicates that both ECR mentees and mentors are acutely aware of these facts, which largely dictate the publishing practices of both ECR mentees and mentors. It is well documented that the traditional metrics ECRs are judged on do not necessarily reflect quality, and they have created a hypercompetitive funding and hiring environment as a consequence of the ‘publish or perish’ academic culture [13, 15–18]. As a further consequence of the pressure induced by the need to publish, this study has shown that a minority of ECR mentees and mentors have experienced authorship disputes. However, positive changes have occurred over the years, with some funding agencies such as the European Molecular Biology Organization (EMBO) choosing to focus more on scientific progress than metrics [92]. Utrecht University [16] and others decided that by early 2022 they would no longer consider impact factor in hiring or promotion decisions. More bold steps such as those taken by EMBO and Utrecht University are required to spur an international culture and policy shift towards fair and equitable recognition of ECR career development.

Another finding of this study is that setting clear expectations, being collegial, mutually respectful, and having regular communication was identified by survey respondents and interviewees as integral features of a healthy ECR mentee-mentor relationship that may affect publishing practices and career development. Communication is commonly identified as an important factor in the initiation and maintenances of mentee-mentor relationships in many professions [93, 94], including those in STEMM [94, 95]. However, as revealed in this study, mentors indicated that they feel unprepared for mentorship and would like training for mentorship. This implied that there are opportunities for academic institutions to focus on faculty development and provide training to ECR mentors so that they may become more effective in their practice. Indeed, the findings of this study also provide valuable information that may assist universities in the development of guidelines surrounding publishing practices, career development, and the formation of postgraduate or postdoctoral charters or program design.

## 5.0 Study limitations

This research adds valuable information to the existing literature regarding ECR mentee-mentor relationship and how this relationship affects ECR publishing practices and career development; however, this study does have notable limitations. First, this study was limited because only approximately a quarter of participants of the survey were ECR mentors (n = 28), which limits the general extrapolation of these findings beyond this cohort to other mentors. Furthermore, it is likely that sampling bias may affect responses due to the respondents own personal interests in this subject area and/or their own values or experiences regarding mentorship. Indeed, overly positive, or negative experiences of mentee-mentor relationship may motivate individuals to response to the survey leading to sampling bias. Response bias may also affect the findings of the survey as it is possible that mentors may not be honest about negative aspects of their own mentorship practices or indeed mentees may not be honest about their contributions and responsibility to establishing and maintaining a healthy mentee-mentor relationship, which is a relationship characterized as a two-way street [96, 97].

Another limitation of this study was the recruitment strategies deployed. Using social media platforms may also affect response rates and induce sampling bias by limiting the study to individuals who take part in social media; however, the advantage of using X is that a wide and varied response can be gathered. Furthermore, social media was used to draw attention to participation in the survey via the polls. However, no demographic information is available to determine what occupation the X respondents were, what level of academia they were part of (if at all), and their career stage was unknown. Furthermore, to see the total results, interested X users need to cast a vote, therefore some votes may be erroneous. Also, with change of leadership at X, it is unclear whether it will be possible to replicate the polls in the future due to a large migration of STEMM researchers away from the platform and changes in operation [44]. Although, it may still be an effective platform to disseminate the survey. The study is also limited by the low number of interview participation; however, the interviews are part of a related ongoing study that is still gathering participants and data at the time of writing this manuscript.

## 6.0 Conclusions

This study has shown that there is varying understanding of what an ECR can defined as. ECR mentees and mentors are largely in agreement with each other regarding publishing practices, although mentees place more value on publishing open access and in prestigious journals for their perceived career impact. Setting clear expectations, being collegial, mutually respectful, and having regular communication are key features of a healthy ECR mentee-mentor relationship that many impact ECR career trajectory. This study adds further understanding to the literature regarding the ECR mentee-mentor relationship and our understanding of how these may affect publishing practices and career progression of ECRs, which may have further implications for policy makers in academia and government.

## Author Contributions statement

Conceptualization, methodology, software, formal analysis, investigation, data curation, writing— original draft preparation, visualization, and project administration, R.L.; writing—review and editing, R.L., M.D., and I.O.D.; Supervision, M.D. and I.O.D.

## Acknowledgements

We would like to thank the teaching team at the Center for Transformative Learning at the University of Limerick, Ireland and colleagues at the Institute for Translational Medicine and Therapeutics, University of Pennsylvania, for their continued support. This research did not receive any funding.

## Ethical Statement

This research has been approved and conducted in accordance with the guidelines and ethical review of the Faculty of Arts, Humanities, and Social Sciences, at the University of Limerick, Ireland (Approval Number 2022-02-10-AHSS). Participants provided informed consent to take part in the survey and interview data presented in this manuscript.

## Data Availability Statement

Survey data is accessible upon request to the corresponding author. Interview data may be embargoed to protect anonymity but may be shared contingent to approval from the University of Limerick Faculty of Arts, Humanities, and Social Sciences Ethics Committee.

## Conflicts of Interest

The authors declare conflicts of interest.

## Notes

### Competing Interest Statement

The authors have declared no competing interest.

