## Supplementary Tables and Figures for "A survey of the mentor-mentee relationship in early career research (ECR): Implications for publishing and career advancement in the STEMM disciplines"

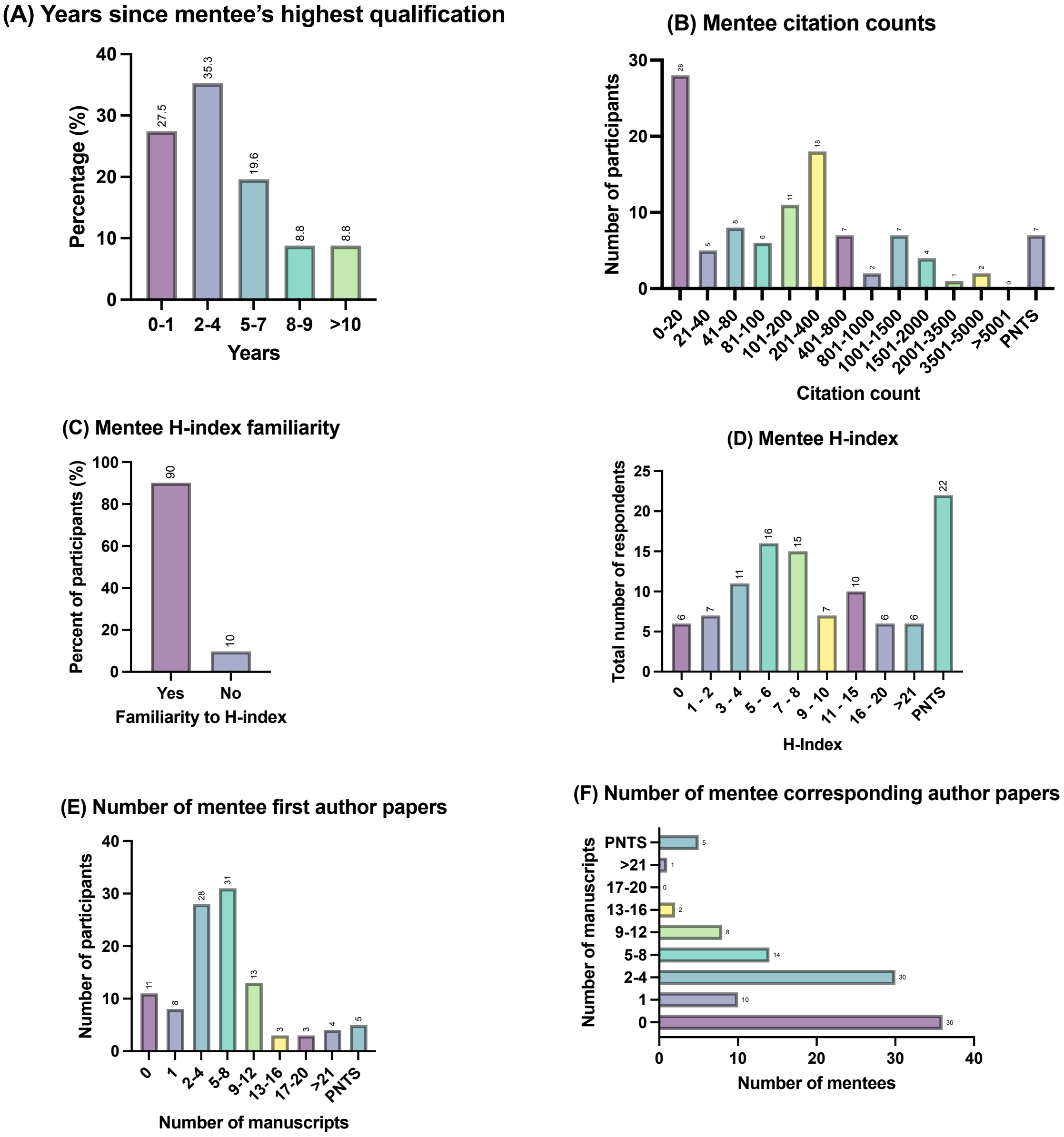


**Figure S1:** ECR mentee responses (n = 106) regarding the so-called traditional academic metrics. (A) The years since the mentee's highest academic qualification; (B) Mentee respondents' indicated citation counts; (C) Mentee's familiarity with the term H-index; (D) Mentee's provided H-index value; (E) Mentee respondents' number of first author publications; (F) Mentee respondents' number of papers where they have been the corresponding/co-corresponding author.

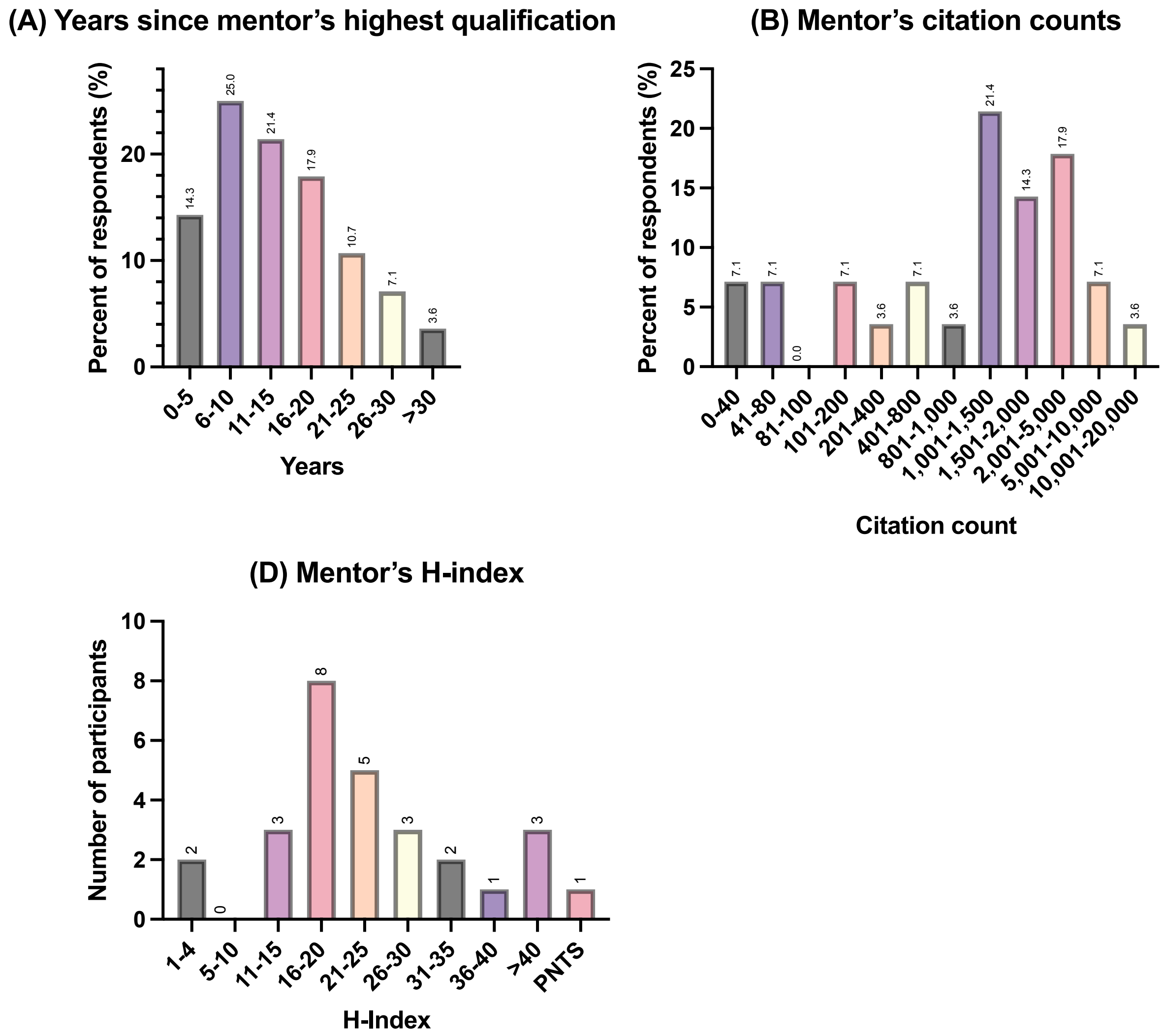


**Figure S2:** ECR mentor responses regarding the so-called traditional academic metrics and years (A) the years since the mentor's highest academic qualification (n = 28); (B) Mentee respondents' (n = 28) indicated citation counts; (C) Mentee's provided H-index value (n = 27).

**Table S1:** Exemplary survey responses to the question addressed to mentees: Are there research journals or publishers that you tend to favor? If so, why? A selection of responses is presented below from a total 79 responses recorded.

| Quote number | Specific responses |
|---|---|
| **1** | *Generally high impact journals.* |
| **2** | *Yes - our field is quite niche, so we tend to publish in only a few select journals from 2 publishers.* |
| **3** | *We tend to favor society journals, as these focus on the audience we are trying to reach, or open access publications in broader journals to increase the visibility of our work.* |
| **4** | *He prefers the CNS journals (Cell, Nature, and Science).* |
| **5** | *Nature journals for the prestige (mentor), and subject specific and open access (mentee)* |
| **6** | *We tend to favor publishers such as Wiley, Springer, and Taylor & Francis as they have a wide range of journals to publish in. Our university recently signed APC agreements with these publishers which is enticing to publish in as the University covers Open Access fees for journals under these publishers.* |
| **7** | *We try to publish in Q1 journals as that is what our university guidelines suggest. Also, we aim for journals with impact factors above 4 if possible.* |
| **8** | *My mentor prefers to aim for the highest impact journal possible for any given project (of course recognizing that some studies are more novel / would appeal to wider audiences / have better chances of being accepted to higher tier journals than others.) I strongly believe that this is not just about prestige, but equally about broader reach and readership.* |
| **9** | *I respect Nature publishing group, Royal Society or ACS journals because of the quality of research work they publish.* |
| **10** | *Open access and society journals.* |
| **11** | *Higher impact journals are favored definitely if the projects research is viewed as more disruptive or groundbreaking to the field because it'll give the readership that view as well and have it be more influential in the field.* |
| **12** | *Nature Science Cell PNAS Impact factor, prestige, popularity.* |
| **13** | *Which journals I choose (and have always chosen even as a grad student because my PhD advisor allowed me a say in the journals) depends on fit, length of time it usually takes to get a paper published in that journal, and past experiences.* |
| **14** | *During my time working with my current mentor, we have primarily sent our publications for review at Cell Press journals for reasons related to scope/readership in one case and invitation for a special issues in another.* |
| **15** | *Yes, those with higher impact, those with positive reputations in terms of research conduct, those with strong content (as distinct from impact).* |

**Table S2:** Exemplary survey responses to the question addressed to mentors: Are there research journals or publishers that you tend to favor? Why/why not? Exemplary responses are presented below from a total 20 responses recorded.

| Quote number | Specific responses |
|---|---|
| **1** | *Yes, reputable, speedy, high impact journals where I know the editors and their priorities.* |
| **2** | *Elsevier and Springer due to the reason of being well established publishing houses.* |
| **3** | *Yes. CNS (Cell, Nature, and Science). Although they are prestigious, they are also legitimate, and peer review is valid.* |
| **4** | *Elsevier- highest impact.* |
| **5** | *Society run journals, smaller open access journals.* |
| **6** | *Yes, high impact Elsevier, Wiley or Nature journals due to their reputation among funders and colleagues.* |
| **7** | *I like open-access journals/publishers, but otherwise, choose first for specialization and then by impact factor/prestige.* |
| **8** | *Impact factor is more important in general.* |
| **9** | *I try to favor Society journals as these give back to community.* |
| **10** | High impact factor journals for the field. Elsevier, Wiley, Nature, or Cell is generally safe. |
| **11** | No, I believe all of them have their specific topic and they are all the same value. |
| **12** | Elsevier and Springer due to the reason of being well established publishing houses. |

**Table S3:** Exemplary survey responses to the question addressed to mentees: Are there research journals or publishers that you or your (supervisor) mentor tend to avoid? If so, why? A selection of responses is presented below from a total 79 responses recorded.

| Quote number | Specific responses |
|---|---|
| **1** | *We try to stick to academic society-run journals where we can so that we can support the field rather than company profits.* |
| **2** | *MPDI and most Frontiers journals, because of their reputation as predatory publishers trying to maximize publication fees rather than facilitating robust peer review.* |
| **3** | *I personally rely on impact a lot. There's a set of great journals in my field, but otherwise I tend to ignore smaller less impactful journals.* |
| **4** | *Anything with MDPI because they have a predatory reputation.* |
| **5** | *Not specifically but I have been advised during research training modules to avoid ones that require a big fee and seem too good to be true as they might not be genuine journals.* |
| **6** | *We have not had the best experience as reviewers of Frontiers journals, so would be hesitant to submit there now (although I do have publications in this family of journals). A previous mentor was very against pre-print servers (e.g., BioRXiv), but I think they are a good way to indicate productivity as an ECR while papers can be held up in revision at larger journals.* |
| **7** | *We avoid journals that we do not fit with their scope.* |
| **8** | *I don't like Elsevier run journals because of the high profit without giving much back to the community.* |
| **9** | *I avoid predatory journals (Not indexed in Web of Science or Scopus database)* |
| **10** | *Yes, predatory journals or journals they have had bad experiences with.* |
| **11** | *Journals outside Q1, because they are not as well considered for the CV.* |
| **12** | *MDPI and Frontiers. Even though I have previously published with both frequently and they have high impact factor journals, their reputation is getting worse, and I don't want that to overshadow my research and CV. My mentor will never and has never published in MDPI.* |

**Table S4:** Exemplary survey responses to the question addressed to mentors: Are there research journals or publishers that you tend to avoid? Why/why not? A selection of responses is presented below from a total 20 responses recorded.

| Quote number | Specific responses |
|---|---|
| **1** | *I try to avoid publishers that have a bad reputation in my field.* |
| **2** | *MDPI because it is predatory, Frontiers because it is expensive and other journals.* |
| **3** | *Low quality, semi predatory journals, and those that don't offer open access.* |
| **4** | *Yes, if I had to pay for publication, I try to avoid publishing a paper on that journal.* |
| **5** | *MDPI - pay to publish (but have published there).* |
| **6** | *Predatory journals/publishers that are aggressive and low-quality.* |
| **7** | *Yes, many of the predatory journals, some are difficult to avoid completely but I prefer an alternative.* |
| **8** | *The ones with high publishing fees, because I have no for that.* |
| **9** | *MDPI, Frontiers, and journals that have high APC. I don't like to support journals that offer open access at high prices and don't offer the subscription model too. It is exclusionary. I have a young lab, we are lucky to be able to support ourselves, we don't need to pay thousands to a publisher.* |
| **10** | *Generally, lately we are avoiding some of the "mega journals".* |
| **11** | *MDPI and Sage... low quality peer review and bad reputations* |
| **12** | *I dislike Bentham. The reason is that they do not supply by free a copy of the final manuscript to authors.* |
| **13** | *The ones with high publishing fees, because I have no funds for that.* |

**Table S5:** Mentees were asked to indicate to what extent they agree with the following statements regarding ECR mentee-mentor relationships and career development (n = 100). Results are expressed as a percent of the total responses provided.

| | **Percent of respondents (%)** | | | | |
|---|---|---|---|---|---|
| **Statements** | **Strongly disagree** | **Somewhat disagree** | **Neither agree nor disagree** | **Somewhat agree** | **Strongly agree** |
| I think that my mentor values my work contributions and opinions | 1 | 3 | 5 | 32 | 59 |
| My mentor encourages and supports my career development | 2 | 7 | 6 | 30 | 55 |
| My mentor encourages me to be collegial and establish my own network | 5 | 4 | 16 | 25 | 50 |
| My mentor has nominated me for grants and awards | 24 | 15 | 23 | 19 | 19 |
| My mentor and I have a collegial relationship | 2 | 4 | 7 | 37 | 50 |
| I will likely collaborate with my current mentor in the future | 6 | 6 | 11 | 25 | 52 |

**Table S6:** In the survey, mentees and mentors were asked to indicate to what extent they agree with the following statements regarding ECR career development? Results are expressed as a percent of the total responses provided by either mentees (n = 94) or mentors (n = 24).

| | **Percent of respondents (%)** | | | | |
|---|---|---|---|---|---|
| **(A) Mentee statements** | **Strongly disagree** | **Somewhat disagree** | **Neither agree nor disagree** | **Somewhat agree** | **Strongly agree** |
| The amount I publish matters for my career development | 0.0 | 1.1 | 3.2 | 26.6 | 69.2 |
| The quality of my published materials matters for my career development | 0.0 | 0.0 | 0.0 | 22.3 | 77.7 |
| H-index is an important factor in my career development | 1.1 | 4.3 | 21.3 | 46.8 | 26.6 |
| Citation counts matter to my career development | 0.0 | 2.1 | 13.8 | 42.6 | 41.5 |
| I have a writing or publication plan | 9.6 | 9.6 | 17.0 | 38.3 | 25.5 |
| I have a career development plan | 7.5 | 9.6 | 17.0 | 38.3 | 25.5 |
| I think future potential employers will focus on publishing metrics such as publication count and h-index when hiring | 2.1 | 7.5 | 20.2 | 36.2 | 34.0 |
| My mentor is invested in my career development | 7.5 | 4.3 | 19.2 | 31.9 | 37.2 |
| | **Percent of respondents (%)** | | | | |
| **(B) Mentor statements** | **Strongly disagree** | **Somewhat disagree** | **Neither agree nor disagree** | **Somewhat agree** | **Strongly agree** |
| The amount an ECR publishes matters for their career development | 0.0 | 8.3 | 0.0 | 20.8 | 70.8 |
| The quality of the published materials is the most important factor for the career development ECRs | 0.0 | 0.0 | 4.2 | 25.0 | 70.8 |
| H-index is an important factor in the career development of ECRs | 4.2 | 16.7 | 20.8 | 45.8 | 12.5 |
| Citation counts matter for the career development of ECRs | 0.0 | 20.8 | 20.8 | 41.7 | 16.7 |
| My institution offers promotions and pathways to tenure for ECRs | 33.3 | 25.0 | 29.2 | 8.3 | 4.2 |
| My institution provides internal grants to promote independence of ECRs | 37.5 | 8.3 | 29.2 | 16.6 | 8.3 |
| I encourage my ECRs to have a career development plan | 0.0 | 0.0 | 12.5 | 25.0 | 62.5 |

**Table S7** Exemplary survey responses to the question addressed to mentors: How does your role as a mentor of ECRs meet your original expectations? A total of 21 written responses were recorded.

| Quote number | Specific responses |
|---|---|
| **1** | *Mentoring ECRs is highly rewarding for my personal satisfaction of the job, but may interfere with some productivity goals, especially if mentoring ECRs falls outside of my own lab personnel.* |
| **2** | *It is harder than it seems. It's difficult to get good grants and good students.* |
| **3** | *Somewhat more challenging. There is no training to be a mentor, you either have it or you don't.* |
| **4** | *More difficult than I thought, but also more rewarding.* |
| **5** | *Wish I had more time to devote to their mentoring and development as well as more resources to maintain them.* |
| **6** | *As I had fast track transformation from postdoc to tenure professorship, I believe that everything I was missing during my ECR stage I am now trying to give to my ECRs.* |
| **7** | *I find it more challenging than I thought. It is difficult to juggle the needs of my students and postdocs with covering administrative work.* |
| **8** | *It is difficult to find the time, and I have few staff.* |
| **9** | *It is as I expected it, with the difference that I have very little time due to heavy teaching load.* |

**Table S8:** Exemplary responses to the question addressed to mentors: Is there anything you would like to add which has not been covered in this survey presented. In total, 9 responses were recorded that included text, which are presented below.

| Quote number | Specific responses |
|---|---|
| **1** | *Goals and expectations should be clear between mentors and mentees.* |
| **2** | *With increasing competition from industry and deceased quality and permanence of research positions tied to certain grants & not institutes themselves, ECRs are seeking and finding other options. Many are leaving academic positions early, which leaves a lot of pressure on mentor to deliver on the project.* |
| **3** | *Mentoring is not part of current professional development, including workloads or progression.* |
| **4** | *I think we all want to do better for ECRs. It is very difficult times, and we have limited time and resources to do a lot of the things for ECRs than we would like. To an extent the ECRs need to manage their PIs too; and make sure that they communicate their needs clearly - prioritizing is really hard and we are so time poor.* |
| **5** | *I feel that there needs to be training not only for the ECRs but also for mentors.* |

**Table S9:** Exemplary quotes regarding publishing and authorship practices from some of the ECR mentees (n = 4) and mentor (n = 2) interviewees.

| Mentee Interviewee number | Excerpts |
|---|---|
| 1 | ***Excerpt regarding publishing:*** *My experience back in country X was not a pleasant one. We had to look for journals APC's. Basically, because we weren't funded that much. So that was quite a hindrance to find journals that we fit the scope of that do not require payment.* |
| 2 | ***Excerpts regarding authorship disputes:*** *In my last the lab where I did my first postdoc, was where things got real. I guess I saw the other side of how favoritism can work. How if you don't push hard enough and you're not the loudest one in the room you can get walked all over and people will steal things [authorships] from you.*<br><br>***Excerpts regarding authorship disputes:*** *I had one project I was working on that went a different route and we needed some help from another postdoc in the lab. That postdoc went ahead and behind my back with the with the PI and came up with a whole set of experiments to completely redirect the project. I was doing these other experiments, you know, but they had set up this whole other side of the project that ended up being the dominant story. We were Co first authors on the paper, but then that became a whole thing. It was my project originally, but then whose name goes first. That became a fight, and it was exhausting. I was like, I don't even care. I just wanted to be a co-first author, but the other girl she fought for being the first-first author [first in appearance]. You don't think it matters, but it does. It does matter when you're applying for things in the future.* |
| 3 | ***Excerpts regarding authorship disputes:*** *I've heard of people being disappointed who contributed to work, who helped kind of manage things and do some practical work and didn't get on the publication. They were disappointed.* |
| 5 | ***Excerpts regarding publishing:*** *When you aim for high impact factor it is assumed that you will have a broad audience… I feel like a lot of people read Nature, Science, and Cell, those three would be like key targets for your paper* |
| **Mentor Interviewee number** | **Excerpts** |
| 1 | ***Excerpts regarding authorship:*** *There have been some disputes, frictions and you know who is going to be the first author or the second author? But I'm trying to minimize the frictions as early as possible. Because if you if you deal with them at the initial stage of the friction, the friction is small and then can be resolved easily. But if you leave it to drag down the road then it becomes a problem. So yeah, I'm trying to be up front with my coauthors saying that OK guys, you can put your names as equal contributors or one of you is first author on this paper and then the 2nd paper, the other one is going to be the first author. You know things like that. Just to keep a balance.*<br><br>***Excerpt regarding publishing:*** *When I choose a journal, I choose it in terms of impact factor and but also speed, speed of publication, speed of publication is quite important these days.* |
| 2 | ***Excerpts regarding the theme of authorship:*** *Not in my group, but yes, I would have. Multiple times or even like, you know, seeing things published without my name there. And then I was like, I did this, I was there collecting the data. So yeah, multiple times unfortunately.* |

| | |
|---|---|
| **3** | ***Excerpts regarding authorship disputes:*** *There was once when we had a big workshop and then we published a paper from that and there was one person who asked if two junior colleagues that joined him in the workshop could be on the paper, but they hadn't contributed anything to the workshop and they hadn't really contributed anything to the paper. So, it wasn't really an argument. It was just a little bit tense on my end.* |

**Table S10:** Exemplary excerpts regarding characteristics of mentorship that are favorable or should be avoided provided by some of the ECR mentee (n = 4) and mentor (n = 3) interviewees.

| (A) Mentee Interviewee number | Excerpts |
|---|---|
| 1 | ***Excerpt regarding mentorship characteristics:*** *I think approachability is one of them. Availability to kind of goes with that. I think the converse is bad communication is a characteristic. If a mentor expects a mentee to do something but has not communicated that very explicitly with the mentee, basically there will be a clash at some point, so communication and approachability are two big characteristics. And then I think experience and career stage too. Yeah, I believe, like, approachable mentors who are well advanced in their career stages are well positioned to be able to guide and further develop their mentees.* |
| 2 | ***Excerpt regarding good mentorship and communication:*** *Communication is such a general term but it's pretty important to be able to reach out to your mentor when you want feedback on something. My mentor will get back to you faster than anybody, which always shocks me. But it's you that needs to initiate that at the start. After communication, you need trust.*<br><br>***Excerpt regarding poor mentorship:*** *If they're just never around, they're never on the radar, like you can't track them down or they don't give you any feedback on anything you give them. It might be something to read like an abstract and they can't give you any feedback. If you asked for their opinion on a grant and they don't give feedback? I feel like they don't even know you then.* |
| 3 | ***Excerpt regarding good mentorship and communication:*** *Communication is very important for mentorship, but it is also like important to be open to feedback and ideas that that your people want to pursue. Maybe not every time. That might be a little bit too general, but receptive to ideas that the mentee wants to pursue.* |
| 4 | ***Excerpt regarding poor mentorship and communication:*** *I suppose a lack of communication. Communication is important to be a good mentor. So, if you're not getting that kind of direct communication with somebody or your kind of leaving them to their own devices, too much. Checking in with them often it is important. Maybe not relying on them [mentors] to schedule the meetings, try to kind of have things, maybe a bit structured.* |
| **(B) Mentor Interviewee number** | **Excerpts** |
| 1 | ***Excerpt regarding the benefits of mentorship:*** *This is similar question to: Is there a benefit for being a teacher? I feel there is. I would think so. There should be benefits. I mean you build a personal relationship with this person by trying to help him or her with his or her career. Then sometimes the comments exchanged during discussion and the points that they come across; I think they can be helpful to the mentor as well. But as I said, it needs to be. It needs to be a very open and transparent relationship between the two of them.*<br><br>***Excerpt regarding poor mentorship:*** *Yeah, I think bad mentors are usually people that think that their research wise is the best and there's nothing else out there.*<br><br>*The other thing is a lack of empathy makes a bad mentor. Also, I would think a mentor needs to be realistic and have both feet on the ground. Some mentors believe that this is so easy. This is what I've done, so if you do exactly the same thing, I've done you'll be successful, but you know every person is unique and the mentor needs to be flexible. If mentors are rigid in their approach. I think this is a bad mentor.* |

| | |
|---|---|
| 2 | ***Excerpt regarding good mentorship:*** *So, I think a good mentor is and is the one who doesn't spend too much time micromanaging because that's what I feel like I'm doing in my mentees research. Another is providing opportunities for their students, and this is again something that I will do and that I'm doing all the time. I'm always suggesting to them to go for conferences to go for training. It's not just the research project that they need to focus. I think they need to be given an opportunity to develop all sorts of skills that are relevant in academia, teaching, supervising, but also presenting and more technical skills like how to analyze your data, whether we're talking about stats, or whether we're talking about qualitative data. So, I think they should be given these opportunities. The thing is that they also need to be on their own a little bit more. They are like your children. You let them fail so that they succeed.*<br><br>*The other thing is that you, you know, you need to make them understand that it is their own projects and you're helping them to achieve their goals and to answer the research questions, but it's their project. And this is the hard thing that I personally find very hard to do.*<br><br>*I have asked our department to organize workshops or even courses on how to supervise students because I feel like the way that I'm supervising students is not sustainable for me in in the long run.*<br><br>***Excerpt regarding poor mentorship and communication:*** *A bad mentor would be a mentor who wouldn't respect the opinion of the student and somebody who would not provide them opportunities for training, for conferences, and somebody who wouldn't be satisfied with whatever they did. Focusing on the negative.* |
| 3 | ***Excerpt regarding mentorship:*** *A characteristic of a good mentor is not being exploitive. There are far too many exploitive people… Another key challenge of being a mentor is instilling the good sound kind of stuff, like integrity, and I find that one really hard to reconcile actually. On some level I could potentially be encouraging my people to become sort of like paper mills and publish all sorts of rubbish, chasing top journals, and doing other things like trying to slice the research up [meaning distribute data across multiple manuscripts rather than one large manuscript to increase the number of publications] and do all of this other stuff. And that's not really going to help much. It might help ECRs get their next job, but it's not sort of like helping them progress in a quality over quantity argument to be made in research. But I think also that some of our quality indicators are not as good as they could be either.* |

**Table S11:** Questions formed for the semi-structured interview for (A) mentees and (B) mentors.

**(A) Mentee Questions**

1. Can you briefly introduce yourself and your background.
2. What is the structure of your research group?
3. Can you tell me about your publishing experiences as an early career researcher (ECR)?
4. What are the opportunities and challenges to being an ECR?
5. What is your experience with working with your supervisor/mentor in regard to publication?
6. What characteristics do you think makes a good or a bad ECR mentor?
7. When working with your supervisor/mentor, what is your process for initiating research projects?
8. When a project is complete, what factors influence the decision to publish your findings?
9. What factors influence who writes and reviews your manuscripts in preparation for publication?
10. What factors influence your route of publication? (e.g., journal selection etc.)
11. What contributions do you think are necessary to be considered for authorship?
12. Who responds to the reviewer comments?
13. Have you ever had disputes about publications or authorship?
14. How does your mentor support your career development?
15. Do you think publication metrics are important for your career development? If so, which ones?
16. If there was advice you could offer other ECRs regarding publishing or career progression, what would it be?
17. If there was advice you could offer ECR mentors regarding mentoring ECRs or supporting career progression, what would it be?
18. Is there anything in your own practice you would change?

**(B) Mentor Questions**

1. Can you briefly introduce yourself and your background.
2. What is the structure of your research group?
3. What does your role as an ECR mentor entail?
4. Are there opportunities or challenges to being an ECR mentor?
5. What characteristics do you think makes a good or bad ECR mentor?
6. When working with ECRs, what is your process for initiating research projects?
7. When a project is complete, what factors influence the decision to publish your findings?
8. What factors influence who writes and reviews your manuscripts in preparation for publication?
9. What factors influence your route of publication? (e.g., journal selection etc.)
10. What contributions do you think are necessary to be considered for authorship?
11. What is your usual process to address reviewer comments?
12. Have you ever had disputes about publications or authorship with ECRs?

13. Can you describe how you contribute to your ECRs career development?
14. Do you think publication metrics are important for ECRs career development?
15. If there was advice you could offer ECRs regarding publishing or career progression, what would it be?
16. If there was advice you could offer other ECR mentors regarding mentoring ECRs or supporting career progression, what would it be?
17. Is there anything in your own practice you would change?
